# 14-3-3ε inhibits premature centriole disengagement by inhibiting the activity of Plk1 and Separase

**DOI:** 10.1101/2024.12.11.627905

**Authors:** Monika A. Jaiswal, Akshay Karn, Aparna Das, Anisha Kumari, Shilu Tiwari, Sorab N. Dalal

## Abstract

The 14-3-3 protein family regulates several pathways in mammalian cells, including centrosome duplication. However, the precise mechanisms by which 14-3-3 paralogs regulate the centrosome cycle remain unclear. To identify the mechanisms by which 14-3-3ε regulates centrosome duplication, we altered two conserved acidic residues in the 14-3-3ε phospho-peptide-binding pocket that regulate complex formation and dissociation with the associated ligands, D127 and E134, to Alanine. Altering these residues to Alanine led to opposing effects on centrosome duplication; the D127A mutant inhibited centrosome duplication, while cells expressing the E134A mutant showed the presence of supernumerary centrosomes. We demonstrate that 14-3-3ε does not inhibit centriole duplication, as reported for 14-3-3γ, but inhibits centriole disengagement. Using a combination of pharmacological and genetic approaches, we demonstrate that 14-3-3ε inhibits the activity of Plk1 and Separase, leading to disengagement defects that ultimately lead to decreased proliferation and cell death. Our work demonstrates that different 14-3-3 paralogs regulate different steps in the centrosome cycle, and disrupting complex formation between 14-3-3ε and Plk1 or Separase could be a novel therapeutic strategy in tumor cells.

## INTRODUCTION

The accurate segregation of the mammalian genome relies on the formation of the mitotic spindle (Reber & Hyman, 2015). Nucleation of the microtubules to form a spindle is dependent on the centrosome, which consists of two centrioles, a mother centriole inherited from the previous cycle and a daughter centriole synthesized in the current cycle, surrounded by the pericentriolar matrix (PCM) (Bornens, 2002; Brinkley *et al*, 1981; Mahen & Venkitaraman, 2012). Centriole duplication is initiated synchronously with DNA replication in S-phase and is limited to once per cell cycle (Hinchcliffe & Sluder, 2001). Before mitosis, the two centrosomes migrate to the two poles, nucleating a bipolar mitotic spindle (Heald & Khodjakov, 2015; Reber & Hyman, 2015). Centrosome amplification or a decrease in centrosome number leads to chromosomal instability and aneuploidy, which are deleterious to the cell (Basto *et al*, 2008; Castellanos *et al*, 2008; Levine *et al*, 2017; Pihan, 2013; Sir *et al*, 2013; Wong *et al*, 2015).

Centriole disengagement is initiated at the end of mitosis and the beginning of G1. Disengagement requires the activity of the Polo-Like Kinase 1 (Plk1) and Separase induced degradation of the S-M linker (Karki *et al*, 2017; Schockel *et al*, 2011; Tsou & Stearns, 2006; Tsou *et al*, 2009). Disengagement is a licensing event for centrosome duplication, as centrioles do not duplicate unless they are ∼80 nm apart (Shukla *et al*, 2015; Tsou *et al*., 2009). During S-phase, the activity of Polo-Like Kinase 4 (Plk4) and cdk2 promotes procentriole biogenesis (Habedanck *et al*, 2005; Kleylein-sohn *et al*, 2007; Meraldi *et al*, 1999). Over-expression of either cdk2 or Plk4 leads to centrosome amplification in mammalian cells. Pro-centriole elongation and the maturation of the daughter centriole from the previous cycle into a mother centriole occurs during G2. The new mother centriole acquires distal and sub-distal appendages and the ability to nucleate PCM (Adon *et al*, 2010; Hinchcliffe & Sluder, 2001; Kleylein-sohn *et al*., 2007; Ohta *et al*, 2018; Whalley *et al*, 2015). Before mitosis, the G1-G2 tether holding the two centrosomes together is degraded in a Plk1 and Nek2A dependent manner (Bahe *et al*, 2005; Fry, 2002; Mardin *et al*, 2011; Mardin *et al*, 2010; Yang *et al*, 2006a) resulting in centrosome separation and formation of a bipolar spindle (reviewed in (Nigg & Stearns, 2011)).

The 14-3-3 protein family has seven paralogs in mammalian cells (reviewed in (Aitken, 2006)). All 14-3-3 paralogs share a similar structure, consisting of nine α-helices that form the monomer, with each monomer in the dimer binding to a phosphorylated peptide via mode I, mode II, or mode III consensus sequences (Coblitz *et al*, 2005; Muslin *et al*, 1996; Rittinger *et al*, 1999; Yaffe *et al*, 1997). The dimerization domain lies in the N-terminus (Liu *et al*, 1995; Xiao *et al*, 1995) and is required for binding to protein ligands (Brunet *et al*, 2002; Shen *et al*, 2003; Tzivion *et al*, 1998). Binding of 14-3-3 proteins to their ligands affects their localization, conformation, stability and function (Dalal *et al*, 1999; Kumagai & Dunphy, 1999; Moorhead *et al*, 1996; Obsil *et al*, 2001; Peng *et al*, 1997; Yaffe, 2002). This could be due to the observation that they function as molecular chaperones preventing aggregation and phase separation (Segal *et al*, 2023). However, it is still not clear how the loss of individual 14-3-3 paralogs leads to the phenotypes observed in mammalian cells (Dalal *et al*, 2004; Sehgal *et al*, 2014; Tilwani *et al*, 2021; Toyo-oka *et al*, 2003) despite some data that suggest that most 14-3-3 paralogs bind to the same set of ligands with different affinities (Segal *et al*., 2023). Previous work from our laboratory has demonstrated that two conserved acidic amino acid residues in the phospho-peptide binding pocket of the 14-3-3 protein family, an Aspartic acid residue and a Glutamic acid residue, regulate complex formation and dissociation from the associated protein ligand (Bose *et al*, 2021; Modi *et al*, 2020). Altering these residues to Alanine increased or decreased binding to the ligand, suggesting that these mutants might help identify specific ligands for individual 14-3-3 paralogs.

Previous work has demonstrated that 14-3-3γ and 14-3-3ε co-purify with the centrosomal fraction (Andersen *et al*, 2003; Pietromonaco *et al*, 1996), and loss of 14-3-3γ and 14-3-3ε leads to centrosome amplification in multiple cell lines due to the premature activation of cdc25C during interphase, leading to premature cdk1 activation and the increased phosphorylation of NPM1 at T199, which results in centrosome amplification (Mukhopadhyay *et al*, 2016). In addition, both these proteins bind to various centrosomal proteins (Mukhopadhyay *et al*., 2016), including Centrin2 and the association of the 14-3-3 proteins with Centrin2 is required for the localization of Centrin2 to the centrosome (Bose & Dalal, 2019a). Loss of 14-3-3γ and 14-3-3ε in HaCaT cells leads to centrosome amplification, but the loss of 14-3-3γ results in clustered mitoses, while loss of 14-3-3ε leads to multi-polar mitoses (Tilwani *et al*., 2021). These results suggested that the two paralogs regulated different proteins required for centrosome licensing and duplication. Indeed, 14-3-3γ prevents centriole duplication by binding to and inhibiting NPM1 phosphorylation by CDK2 (Bose *et al*., 2021). In contrast, the results in this report suggest that 14-3-3ε inhibits centriole disengagement, thereby preventing centrosome licensing. Using genetic and pharmacological approaches, we demonstrate that 14-3-3ε inhibits the function of Plk1 and Separase, thereby inhibiting centriole disengagement. A mutant of 14-3-3ε that shows decreased binding to Plk1 and Separase shows premature disengagement during interphase. Our results identify a novel mechanism by which 14-3-3ε regulates centriole disengagement and demonstrate that different 14-3-3 paralogs affect different steps in the centrosome duplication pathway.

## RESULTS

### 14-3-3ε inhibits centriole disengagement

Loss of two 14-3-3 paralogs, 14-3-3ε and 14-3-3γ, leads to centrosome amplification in multiple cell types (Bose *et al*., 2021; Mukhopadhyay *et al*., 2016; Tilwani *et al*., 2021). However, the loss of these two paralogs resulted in different mitotic outcomes in cells, suggesting that they affected different components in the centrosome duplication pathway. As the loss of 14-3-3ε also led to centrosome amplification, we wished to determine the mechanisms by which 14-3-3ε affected centrosome duplication. As a first step, we altered the conserved Aspartic and Glutamic acid residues in 14-3-3ε to Alanine, D127A and E134A, respectively, and determined their ability to affect centrosome duplication. We transfected the mOrange tagged 14-3-3εWT and mutant constructs (D127A, E134A and D127AE134A) into HCT116 cells, blocked the cells in mitosis with nocodazole and stained the cells with antibodies to pericentrin to determine centrosome number. A plasmid expressing mOrange served as a vector control. As shown in Fig1A-B, most of the cells transfected with WT 14-3-3ε showed two centrosomes as did the vector control. In contrast, a significant number of cells transfected with the D127A mutant or the E134A mutant showed a single centrosome or centrosome amplification, respectively (Fig1A-B), a phenotype similar to that observed with similar mutants in 14-3-3γ (Bose *et al*., 2021). The 14-3-3γ double mutant showed a phenotype similar to WT 14-3-3γ (Bose *et al*., 2021). However, the 14-3-3ε D127AE134A mutant showed a phenotype similar to the D127A mutant (Fig1A-B). Similar results were observed in multiple cell lines (Fig1C-E and EV Fig 1A-C). All the mutants were expressed at equivalent levels in the different cell lines (EVFig1D). To confirm that these results were due to changes in 14-3-3ε, we generated 14-3-3ε knockout lines (14-3-3ε KO) using Crispr/Cas9 in HCT116 and RPE1 hTERT cells and expressed either WT 14-3-3ε or the mutant constructs in these cells. Loss of 14-3-3ε led to increased centrosome amplification in HCT116 and RPE1 hTERT cells compared to the parental cells. Restoration of WT 14-3-3ε reversed the phenotype, and a further decrease in centrosome amplification was observed upon expression of the D127A mutant or the D127AE134A mutant. In contrast, the E134A mutant did not show a further increase in centrosome amplification in these cells (Fig1F-G and EVFig1E-F). All mutants were expressed at equivalent levels in these cells (EVFig1G-H). The loss of another paralog, 14-3-3ζ, did not alter the centrosome number (EVFig1I-K), suggesting that the phenotypes observed are due to the loss of 14-3-3ε.

**Figure 1.**
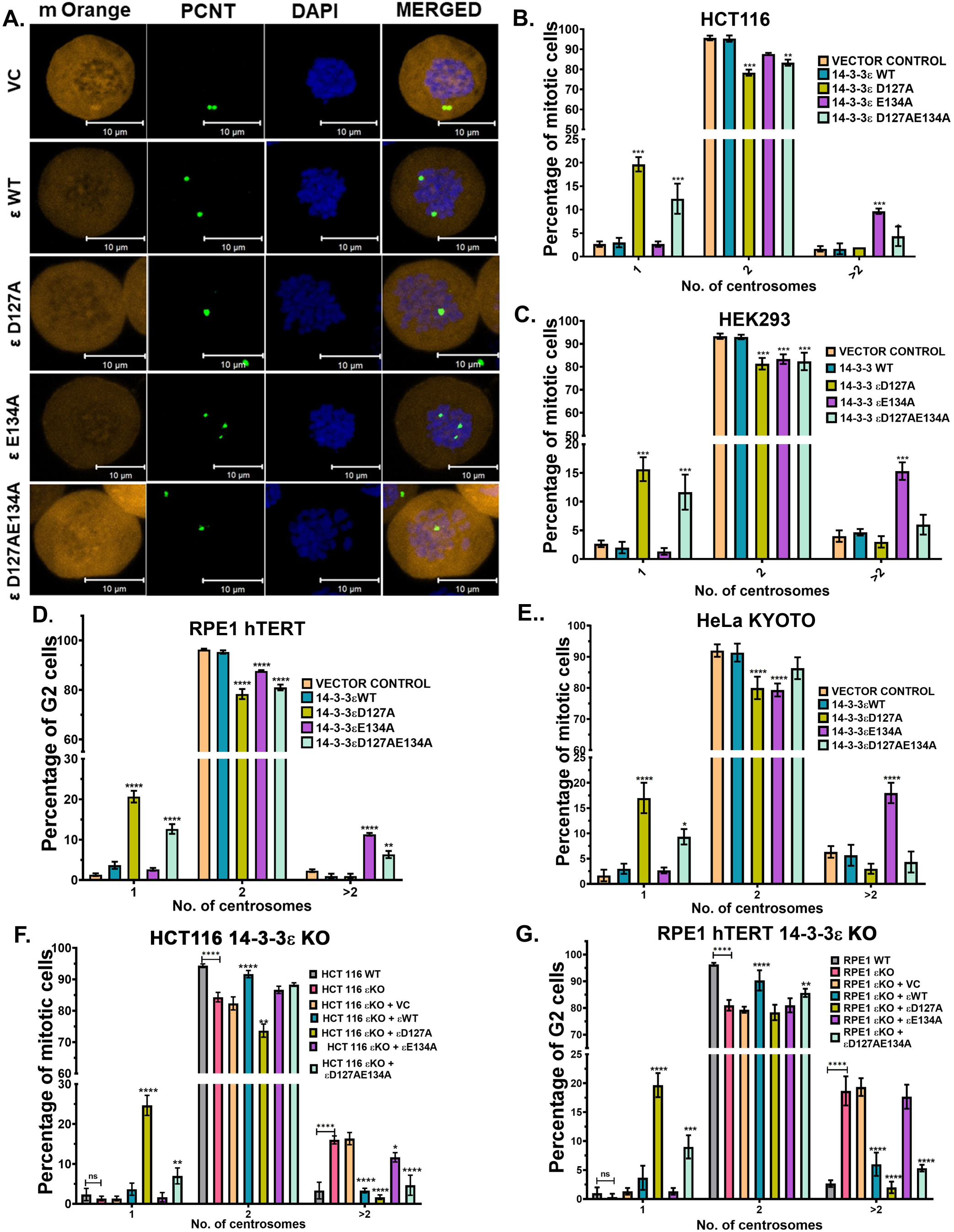
14-3-3ε mutants alter centrosome numbers in different cell lines. **(A- B)** HCT116 cells were transfected with the mOrange vector control (VC) and mOrange tagged wild type (WT) and mutant (D127A, E134A, D127AE134A)14-3-3ε constructs. The cells were arrested in mitosis with nocodazole and stained with antibodies to percentrin (green) and counterstained with DAPI (blue), and the number of mitotic cells showing 1, 2 or >2 centrosomes was determined in 3 independent experiments. Representative images are shown (A), and the mean and standard deviation of 3 independent experiments are plotted (B). **(C-G)** The number of cells transfected with the mOrange vector control (VC) and mOrange tagged wild type (WT) and mutant (D127A, E134A, D127AE134A)14-3-3ε constructs showing 1, 2 or >2 centrosomes were determined in HEK 293 (C), RPE1-hTERT (D), HELA KYOTO (E), HCT 116 14-3-3εKO (F) and RPE1 hTERT 14-3-3εKO (G). Three cell lines were arrested in prophase (C, E and F), while the RPE1hTERT cell lines (D and G) were arrested in G2 with a cdk1 inhibitor. The mean and standard deviation from three independent experiments is plotted. Note that compared to the VC and WT transfected cells, there is a significant increase in cells with a single centrosome or cells with >2 centrosomes in cells transfected with the D127A and D127AE134A mutants and the E134A mutant, respectively. Where indicated p-values were obtained using 2-way ANOVA (Tukey’s multiple comparison). *p <0.05, **p < 0.01, ***p < 0.001, ****p <0.0001. Scale = 10μm

The phenotypes observed upon expression of the different mutants could be due to defects in disengagement, defects in duplication or separation before mitosis (Fig 2A) (reviewed in (Nigg & Stearns, 2011)). To identify the nature of the defect, HCT116 and HEK293 cells were transfected with the 14-3-3ε constructs and GFP-Centrin2 and mitotic cells were stained with antibodies to pericentrin. All constructs were expressed at similar levels in both cell types (EVFig2A). Most of the vector control and WT 14-3-3ε transfected cells showed two pericentrin foci, each containing two centrin foci as expected (Fig2B-D and EVFig2B). In contrast, many D127A and D127AE134A transfected cells showed one pericentrin foci with two centrin foci (Fig2B-D and EVFig2B). In the E134A transfected cells that showed centrosome amplification, most pericentrin foci were associated with only one centrin focus, suggesting that loss of 14-3-3ε function led to a defect in disengagement with an absence of disengagement occurring in cells transfected with the D127A mutant and premature disengagement in cells transfected with the E134A mutant (Fig2B-D and EVFig2B). This phenotype is distinct from the results we observed with 14-3-3γ, where a defect in centriole duplication was observed (Bose *et al*., 2021).

**Figure 2.**
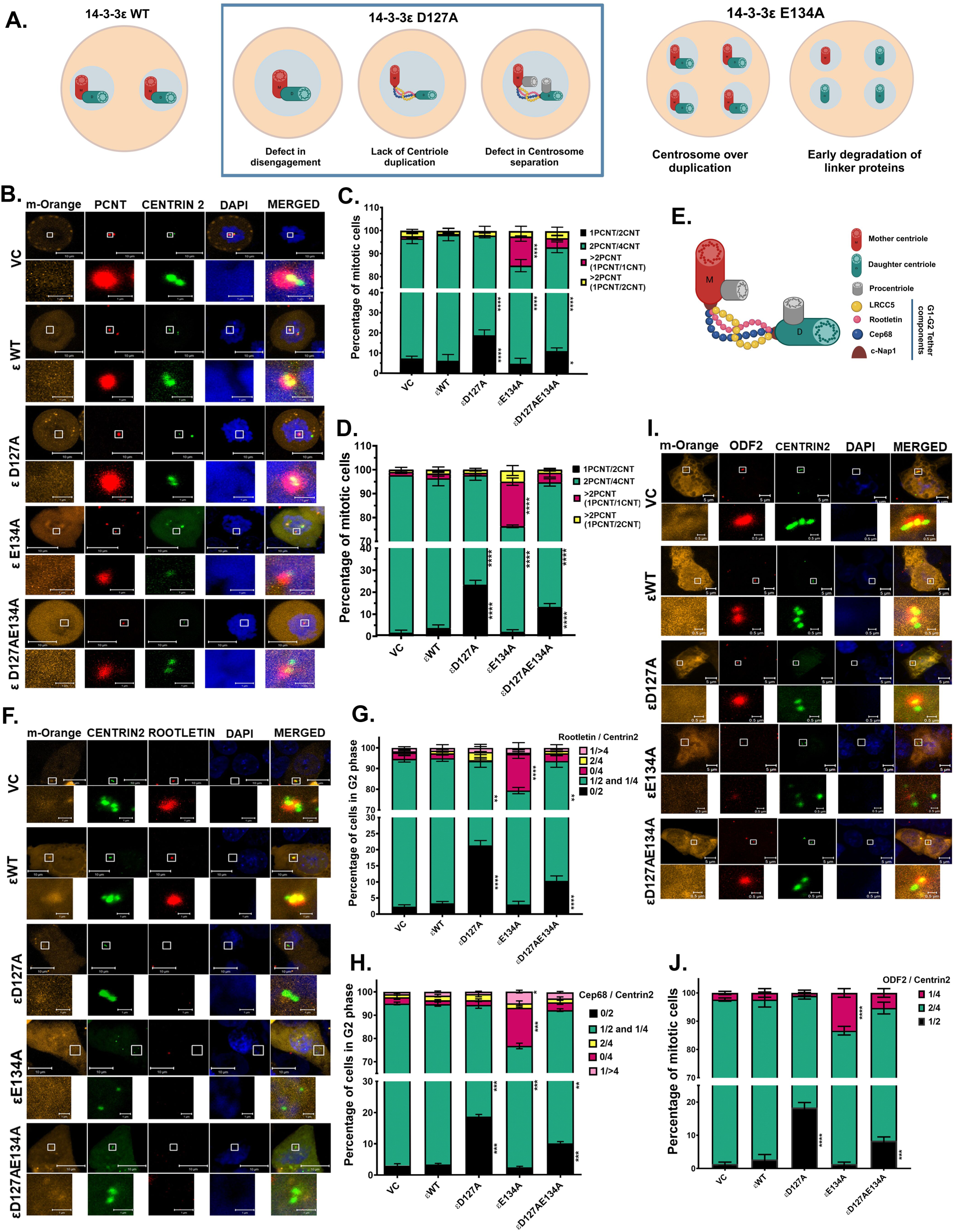
14-3-3ε prevents centriole disengagement. **(A)** Cartoon depicting the potential defect in centrosome duplication in cells expressing 14-3-3ε mutant constructs. **(B-D)** HCT116 and HEK293 cells were co-transfected with GFP Centrin2 (green), and the mOrange tagged WT and mutant 14-3-3ε constructs (D127A, E134A, D127AE134A) (orange), arrested in mitosis with nocodazole, fixed and stained with antibodies to pericentrin (red) and counterstained with DAPI (blue). Representative images for HCT116 cells are shown (B). The pericentrin to centrin (PCNT/CNT) ratio was determined in three independent experiments for HCT116 cells (C) and HEK293 cells (D). Note that cells expressing the D127A and D127AE134A mutants show an increase in cells with a single pericentrin focus with two centrin foci, while cells expressing the E134A mutant show a significant increase in cells with multiple pericentrin foci with a single centrin foci. **(E)** Cartoon depicting the component and localization of G1-G2 tether. **(F-G)** HCT116 cells transfected with the indicated constructs were arrested in G2 with a cdk1 inhibitor, stained with antibodies to the G1-G2 tether protein Rootletin and counterstained with DAPI. Representative images are shown (F), and the Rootletin to Centrin ratio was determined in three independent experiments, and the mean and standard deviation of three independent experiments are plotted (G). Scale = 10μm, inset scale = 1μm. **(H)** HCT116 cells were transfected with the indicated constructs, arrested in G2, stained with antibodies to Cep68 and counterstained with DAPI. The Cep68 to Centrin2 ratio was determined in three independent experiments, and the mean and standard deviation were plotted. **(I-J)** HCT116 cells were transfected with the indicated constructs, arrested in G2 and stained with antibodies to ODF2. Representative images are shown (I), and the mean and standard deviation of three independent experiments are plotted (J). Scale= 5μm, inset scale= 0.5μm. p-values were obtained using 2-way ANOVA (Tukey’s multiple comparison). *p <0.05, **p <0.01, ***p <0.001, ****p <0.0001. Scale = 10μm, inset scale= 1μm

Centriole disengagement requires degradation of the S-M linker at the end of mitosis and the beginning of G1, followed by the formation of the G1-G2 tether and subsequent centriole duplication (Fig2E and (Nigg & Stearns, 2011)). To determine if the G1-G2 tether had formed in these cells, we transfected the 14-3-3ε constructs along with GFP Centrin2 into these cells, arrested them in G2 by treatment with a cdk1 inhibitor and stained with antibodies to different G1-G2 tether proteins. Cells expressing the vector control or WT 14-3-3ε showed that the GFP Centrin2 foci co-localized with staining for the G1-G2 tether markers, Rootletin, and Cep68 (Bahe *et al*., 2005; Graser *et al*, 2007) (Fig2F-H and Fig EV2C). In contrast, in cells expressing the D127A and D127AD134A mutants, a pair of centrin foci did not co-localize with the G1-G2 tether markers, while in cells expressing E134A, the single centrin foci were not localized with the G1-G2 tether markers (Fig2F-H and EVFig2C). However, staining with antibodies to the mother centriole-specific marker, ODF2, which is required to maintain centriole cohesion (Yang *et al*, 2018), showed that a majority of cells transfected with the vector control or WT 14-3-3ε show the presence of two ODF2 foci and four centrin foci as they have two mature mother centrioles. In contrast, a significant percentage of cells transfected with D127A show just one ODF2 foci with two centrin foci and the E134A cells show multiple centrin foci, among which only one centrin focus is associated with an ODF2 focus, suggesting that these are either daughter centrioles that have disengaged from the mother or daughter centrioles from the previous cycle that have not undergone maturation due to premature disengagement from the newly synthesized procentriole (Fig2I-J). All the transfected cells showed equivalent levels of the WT and mutant 14-3-3ε proteins (EV Fig 2D). Taken together, these results suggest that 14-3-3ε inhibits centrosome disengagement.

### The single and multiple centrosomes observed in the 14-3-3ε mutants anchor microtubules but show mitotic defects

Injecting cells with a monoclonal antibody to glutamylated tubulin can result in centriole fragmentation (Abal *et al*, 2005; Bobinnec *et al*, 1998a; Bobinnec *et al*, 1998b). To determine if the centrioles observed in the transfected cells had alterations in the levels of glutamylated tubulin, we performed staining with the antibody to glutamylated tubulin, GT335 (Abal *et al*., 2005; Bobinnec *et al*., 1998a; Bobinnec *et al*., 1998b). The 14-3-3ε proteins are present at equivalent levels (EVFig3A top panel), and the levels of glutamylated tubulin at the centriole in cells transfected with the vector control or the WT or mutant 14-3-3ε constructs did not show significant differences (EVFig3B and Fig3A) suggesting that any defects observed were not due to changes in the levels of glutamylated tubulin at the centrosome. We then generated stable lines expressing the WT and mutant 14-3-3ε constructs in HCT116 cells and measured centriole length and the inner and outer diameter of the centriole using electron microscopy (EVFig3C). Expression of WT 14-3-3ε or the D127A and D127AE134A mutants did not result in a significant change in these phenotypes as compared to the vector control (Fig3B-D). In contrast, the centrioles were significantly shorter in cells expressing the E134A mutant and had a smaller outer diameter than in cells expressing WT 14-3-3ε (Fig3B&D). No change in the inner diameter of the centriole was observed in these cells (Fig3C). These results suggested a defect in the centriole maturation in these cells (reviewed in (Sullenberger *et al*, 2020)). To determine if these altered centrosome phenotypes resulted in a change in proliferation, the m-Orange tagged WT and mutant constructs were transfected into cells, and the number of mOrange-positive cells was monitored over time. The number of mOrange positive cells at each time point was normalized to the initial number of mOrange positive cells present at the start of the experiment for each construct. As shown in Fig3E, cells expressing WT 14-3-3ε showed a higher growth rate than cells expressing the D127A and E134A.

**Figure 3.**
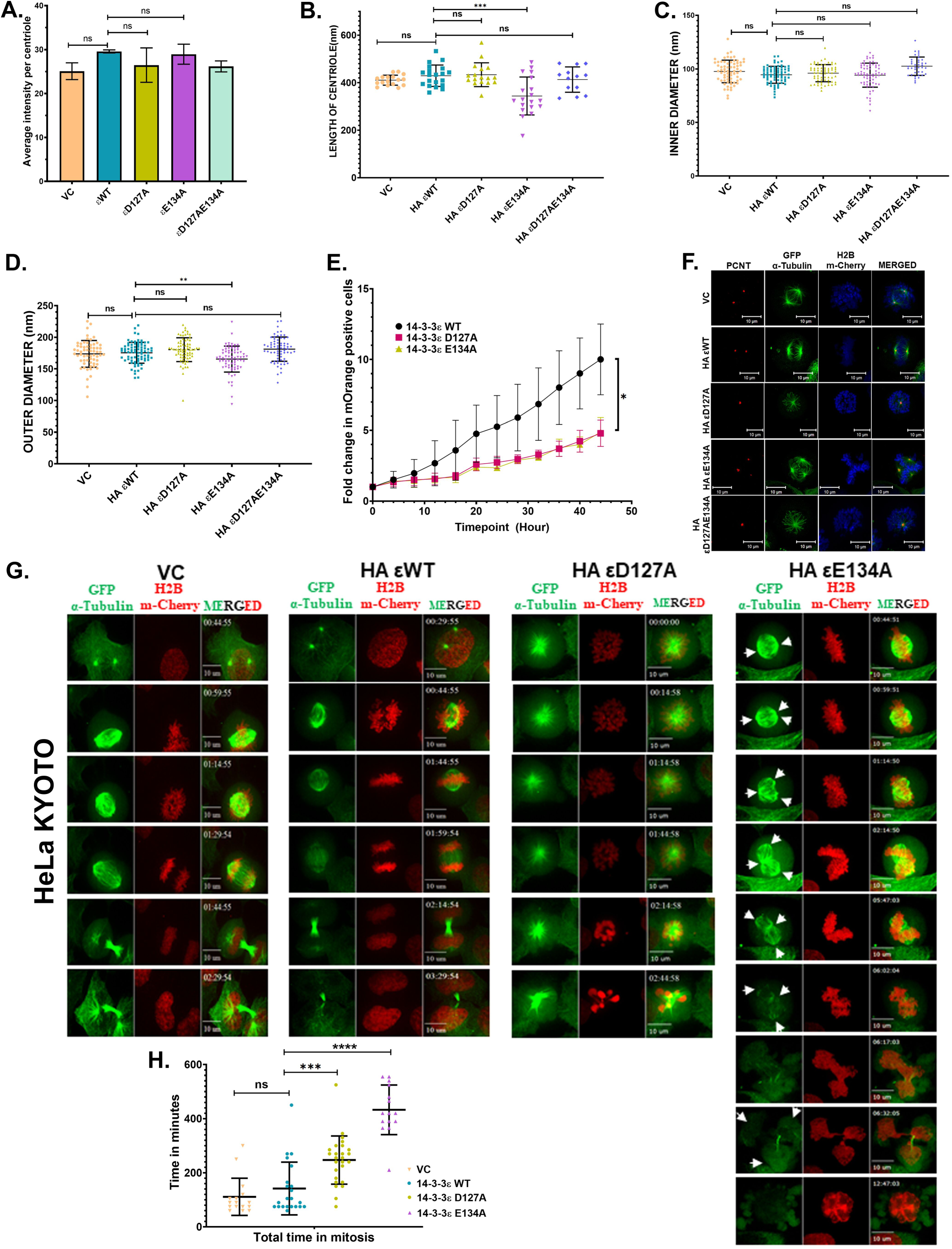
Expression of the 14-3-3ε mutants leads to a mitotic delay and cell death. **(A)** HCT116 cells transfected with the indicated constructs were arrested in mitosis and stained with antibodies to glutamylated tubulin (GT335), and the signal intensity was measured. The mean and standard deviation from three independent experiments are plotted, and p-values are obtained using the student’s unpaired t-test with Welch’s correction. **(B-D)** HCT 116 cells stably expressing the HA vector control (VC) and HA-tagged 14-3-3ε constructs (WT, D127A, E134A and D127AE134A) were synchronized in the G2-M phase and processed for scanning electron microscopy. The length of the centriole (n>15) (B), inner diameter of the centriole (n>39) (C) and outer diameter of the centriole (n>69) (D) were measured in three independent sets and the mean and standard deviation are plotted. **(E)** The proliferation of HCT116 cells expressing mOrange vector control and mOrange tagged 14-3-3ε WT and mOrange tagged 14-3-3ε mutants (D127A and E134A) was measured in an Incucyte live cell analysis system. Images of the same field were acquired every hour and the mean and standard deviation of the normalised counts (mOrange positive/total number of cells) per field from three independent experiments were plotted against time (hours). **(F)** HeLa KYOTO cells stably transfected with HA vector control, and HA-tagged 14-3-3ε constructs (WT, D127A, E134A, D127AE134A) were arrested in mitosis and stained with antibodies against pericentrin (red). The cells express GFP α-tubulin (green) and H2B m-cherry (blue). Scale = 10μm. (G-H) HeLa KYOTO cells stably expressing HA vector control and HA-tagged 14-3-3ε constructs (WT, D127A and E134A) were imaged over 24 hours in an Olympus 3i spinning disc microscope. Representative images of the same cell acquired over the time period for GFP α-tubulin (green) and H2B m-cherry (red) are shown with the time stamp in the upper left corner and scale bar (10μm) in the bottom left corner. The arrowheads indicate MTOCs that show the premature disengagement of centrioles **(G)** The mean and standard deviation of the time spent in mitosis of ≥15 cells across three independent experiments is plotted (H). Note that in contrast to the VC or WT-expressing cells, the D127A cells never complete mitosis, and the E134A cells undergo multipolar mitosis. p-values were obtained using one-way ANOVA (Tukey’s multiple comparison). *p <0.05, **p <0.01, ***p <0.001, ****p <0.0001.

To determine if these centrosomes could anchor microtubules, we transfected the mOrange tagged WT and mutant 14-3-3ε constructs into HCT116 cells and stained them with antibodies to pericentrin and α-tubulin. Cells transfected with the vector control or WT 14-3-3ε showed typical bipolar spindles in mitotic cells. In contrast, cells transfected with the D127A and D127AE134A mutants showed the presence of monopolar spindles, while cells transfected with the E134A mutant showed multipolar spindles (EVFig3D). To further examine the mitotic phenotypes resulting from the expression of these mutants, we stably transfected the WT and mutant 14-3-3ε constructs into HeLa Kyoto cells that express EGFP-tubulin and H2B-mCherry (Bastos & Barr, 2010). The transfected cells were stained with antibodies to pericentrin. Cells transfected with the vector control or WT 14-3-3ε showed typical bipolar spindles in mitotic cells. In contrast, cells transfected with the D127A and D127AE134A mutants showed the presence of monopolar spindles, while cells transfected with the E134A mutant showed multipolar spindles (Fig3F). We then used these stably transfected cells to determine mitotic outcomes in live cell imaging experiments. Cells expressing WT 14-3-3ε completed mitosis in around 80 minutes (Fig3G-H and EV videos 1-2). In contrast, cells expressing the D127A mutant that showed mono-polar spindles never exited prophase and ultimately died, while cells expressing the E134A mutant that showed multi-polar mitoses completed mitosis in around 280 minutes or died due to a prolonged metaphase arrest (Fig3G-H EV videos 3-4). These results are consistent with our observations that cells expressing these mutants show a decrease in proliferation, which is probably because a significant percentage of cells with a single centrosome die in prophase while the cells undergoing multi-polar mitosis also die as previously described (Ganem *et al*, 2009).

### 14-3-3ε inhibits centriole disengagement by inhibiting the activity of Plk1 and Separase

The activity of Polo-Like Kinase 1 (Plk1) and Separase induced degradation of the S-M linker (Karki *et al*., 2017; Schockel *et al*., 2011; Tsou & Stearns, 2006; Tsou *et al*., 2009) are required for centrosome licensing and duplication. As the defect observed in cells transfected with the 14-3-3ε mutants seemed to be a lack of or premature disengagement, we asked if 14-3-3ε formed a complex with Plk1 or Separase. GST pulldown assays demonstrated that GST-14-3-3ε formed a complex with both Plk1 and Separase in contrast to GST alone or the ligand binding defective 14-3-3ε mutant GST-14-3-3εK50E (Brunet *et al*., 2002) (Fig4A and EVFig4A). Similarly, co-immunoprecipitation experiments in HCT116 cells demonstrated that 14-3-3ε formed a complex with both Plk1 and Separase (Fig4B and EVFig4B-C). This was true whether the antibody used for immunoprecipitation was raised against 14-3-3ε, Plk1 or Separase. Proximity ligation assays (PLA) also demonstrated that 14-3-3ε formed a complex with both Plk1 and Separase in cells and cdc25C (Dalal *et al*., 2004) served as a control (Fig4C-D). These results suggested that 14-3-3ε inhibited the function of Plk1 and Separase, thereby preventing premature degradation of the S-M linker. If our hypothesis is correct, one prediction would be that the levels of the S-M linker protein, Cep215, whose degradation occurs before disengagement (Graser *et al*., 2007; Pagan *et al*, 2015), would be altered in cells expressing the different 14-3-3ε mutants. When we stained cells transfected with WT or mutant 14-3-3ε constructs and stained them with antibodies to Cep215, we observed that the intensity of Cep215 staining at the centrosome was highest in cells expressing the D127A mutant as compared to WT 14-3-3ε (Fig4E-F). In contrast, cells expressing E134A showed decreased staining at the centrosome for Cep215 (Fig4E-F). Consistent with these observations, centrosome size as measured by Cep215 staining was elevated in cells expressing the D127A mutant (Fig4G). In contrast, the distance between the two centrioles was elevated in cells expressing the E134A mutant (Fig4H).

**Figure 4.**
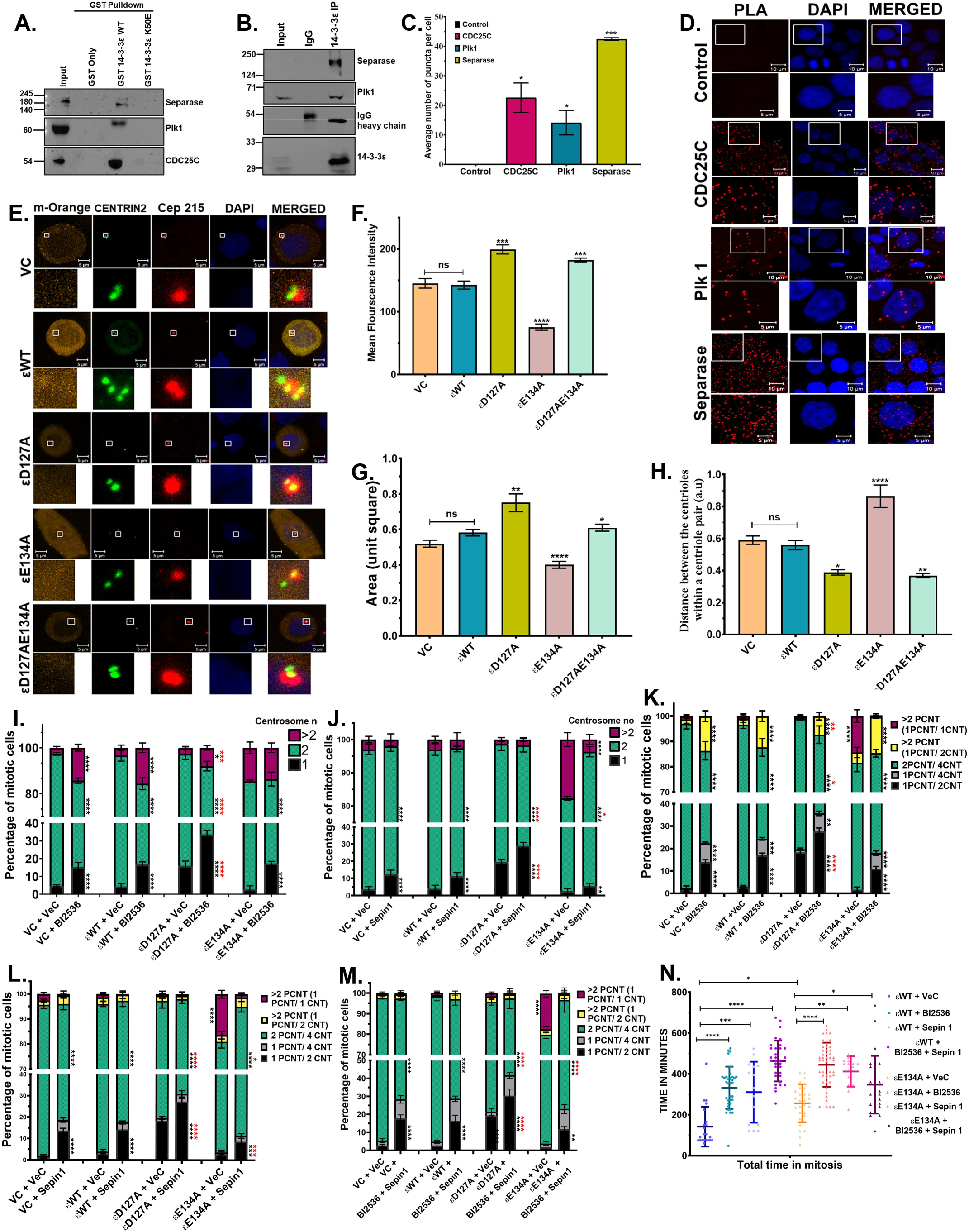
14-3-3ε forms a complex with Plk1 and Separase. **(A)** Protein extracts prepared from HCT116 cells were incubated with the indicated GST fusion proteins, and the reactions resolved on SDS-PAGE gels, followed by Western blotting with the indicated antibodies. Input = 10%. Note that 14-3-3ε forms a complex with Plk1 and Separase in contrast to GST alone or the ligand-binding deficient mutant 14-3-3εK50E. **(B)** Protein extracts prepared from HCT116 cells were incubated with either a non-specific IgG or antibodies to 14-3-3ε, and the reactions resolved on SDS-PAGE gels followed by Western blotting with the indicated antibodies. Input = 10%. **(C-D)** Proximity Ligation Assays (PLA) were performed in HCT116 cells, using antibodies against 14-3-3ε and either Plk1, Separase and cdc25C. Representative images are shown (D), and the mean and standard deviation of the number of puncta per cell from three independent experiments are plotted (C). p-values were obtained using unpaired student’s t-test with Welch’s correction. Scale = 10μm, inset scale = 5μm. **(E-H)** HCT116 cells overexpressing GFP Centrin2 and the mOrange vector control and mOrange tagged 14-3-3ε constructs (WT, D127A, E134A) were arrested in mitosis and stained with antibodies Cep215 and counter stained with DAPI. Representative images are shown (E), and the mean, and standard error from the mean from three independent experiments of the intensity of Cep215 staining (F), centrosome size (G) and distance between the centriole pair (H) pair is plotted. Scale = 5μm and inset scale = 0.5μm. **(I-J)** Cell lines were transfected with the mOrange vector control (VC), and mOrange tagged wild type (WT) and mutant (D127A, E134A, D127AE134A)14-3-3ε constructs and were treated with BI2536 and/or Sepin1 or the vehicle control (DMSO). The mean and standard deviation of the number of centrosomes in mitotic cells from three independent experiments are plotted for cells treated with BI2536 (I) or Sepin-1 (J). **(K-M)** HCT 116 cells were transfected with the mOrange vector control (VC), and mOrange tagged wild type (WT) and mutant (D127A, E134A, D127AE134A)14-3-3ε constructs and GFP Centrin2 and were treated with BI2536, Sepin-1, both BI2356 and Sepin-1 or the vehicle control (DMSO). The mean and standard deviation of the pericentrin to centrin2 ratio from three independent experiments is plotted for BI2536 (K), Sepin-1 (L) and BI2536 and Sepin1 (M). **(N)** HeLa KYOTO cells stably expressing HA 14-3-3ε WT or HA 14-3-3ε E134A were treated with vehicle control (DMSO), BI2536, Sepin-1 or both BI2536 and Sepin-1 and were imaged for 20 hours at intervals of 20 minutes. The mean and standard deviation from ≥ 15 cells across three independent experiments is plotted. p values were obtained using one-way ANOVA (Tukey’s multiple comparison). (*(black) comparison within the group with respective vehicle control, *(red) comparison across the group with 14-3-3ε WT treated with drug) *p <0.05, **p <0.01, ***p <0.001, ****p <0.0001.

As Plk1 and Separase are required for the degradation of the S-M linker we asked whether small molecule inhibitors of Plk1 and Separase, BI2536 (Lenart *et al*, 2007; Steegmaier *et al*, 2007) and Sepin-1 (Zhang *et al*, 2014) respectively, alter the phenotypes observed upon expression of the 14-3-3ε constructs. Treatment of the cells with the inhibitors did not alter the levels of the transfected constructs (EVFig4D). The cell cycle profile of the treated cells is shown in EVFig4E. Treatment of cells with the Plk1 inhibitor BI2536 led to a significant increase in the percentage of cells with a single centrosome compared to the vehicle control (Fig4I). We also observed an increase in the percentage of cells with >2 centrosomes, which might be due to a failure of cytokinesis followed by centrosome duplication in the subsequent cycle as previously reported (Kasahara *et al*, 2013) (Fig4I). Treatment with Sepin-1 resulted in an increase in the percentage of cells with a single centrosome in cells transfected with the 14-3-3ε D127A mutant and a decrease in the percentage of cells showing centrosome amplification in cells transfected with the 14-3-3ε E134A mutant (Fig4J). When we examined the pericentrin to centrin ratio in cells treated with BI2356, we observed that treatment with BI2356 led to a significant decrease in disengagement as the number of cells with only one pericentrin focus and two centrin foci or one pericentrin focus with four pericentrin foci increased significantly, while the number of cells with one pericentrin focus containing a single centrin spot decreased significantly with a concomitant increase in normal centrosome number in the E134A transfected cells (Fig4K and EVFig4F). Similar results were observed when we treated cells with Sepin-1 (Fig4L and EVFig4G). No additive effect was observed when cells were treated with both inhibitors (Fig4M and EVFig4H), which is consistent with the observation that Plk1 activity is required to promote the cleavage of substrates by Separase (Karki *et al*., 2017; Schockel *et al*., 2011; Tsou & Stearns, 2006; Tsou *et al*., 2009). One possible prediction from these experiments is that treatment with the inhibitors would extend the time spent in mitosis. We treated the HeLa lines expressing the WT and E134A mutant with either BI2536 or Sepin-1 or both inhibitors, followed by live cell imaging (EVFig4I-Kand EV videos 5-12). The amount of time spent in mitosis was significantly increased for cells that expressed both WT 14-3-3ε and the E134A mutant of 14-3-3ε (Fig4N). These results suggest that 14-3-3ε might regulate the function of both Plk1 and Separase.

Previous experiments have demonstrated that Plk1 can bind to 14-3-3ζ when phosphorylated at Serine 330 or 597 and 14-3-3γ when phosphorylated at Serine 99 (Du *et al*, 2012; Kasahara *et al*., 2013). As a first step, we determined the interaction of the WT and 14-3-3ε mutants with Plk1. Co-immunoprecipitation and GST pulldown assays demonstrated that the D127A mutant of 14-3-3ε showed a greater interaction with Plk1 than WT 14-3-3ε (Fig5A and EVFig5A). In contrast, the E134A mutant bound to Plk1 at a lower efficiency than WT (Fig5A and EVFig5A). Similar results were observed with cdc25C (EVFig5A). The changes in the association of the 14-3-3ε mutants with Plk1 are not due to a change in their overall structure as determined by their CD spectra (EVFig5B). Similar results were observed in PLA assays, demonstrating that the interaction occurred at the centrosome (Fig5B-C). We then altered a potential 14-3-3 binding site in Plk1 at S99 to Alanine (S99A), Aspartic acid (S99D) and Glutamic acid (S99E) and determined the interaction of these mutants with 14-3-3ε. As shown in Fig5D, co-immunoprecipitation assays demonstrated that the S99A mutant failed to bind to 14-3-3ε, unlike the S99D and S99E mutants, which serve as phospho-mimetic mutants. As controls, we altered another potential 14-3-3 binding site, a Threonine residue at 210 to Alanine (T210A) and demonstrated that this mutant formed a complex with 14-3-3ε (Fig5D). Similar results were obtained in PLA assays (Fig5E and EVFig5D), and all the mutants were expressed at equivalent levels (EVFig5C). We then evaluated the ability of these mutants to affect centrosome number. The expression of the S99A mutant of Plk1 resulted in a significant decrease in the percentage of cells with a single centrosome in cells expressing the 14-3-3ε D127A mutant. In contrast, the phospho-mimetic mutants (S99D and S99E) or the WT Plk1 did not significantly alter the percentage of cells with a single centrosome (Fig5F and EVFig5D). In cells expressing 14-3-3ε E134A, the S99A mutant of Plk1 stimulated a further increase in the percentage of cells showing supernumerary centrosomes as compared to WT Plk1 or S99D and S99E (Fig5F and EVFig5E). To determine the role of these Plk1 mutants in regulating centriole disengagement, the WT or mutant Plk1 constructs were transfected into HCT116 cells, and the pericentrin to centrin ratio was determined in cells arrested in mitosis. The expression of WT Plk1 led to a significant increase in centrosome amplification compared to the vector control (Fig5G). In contrast, the 14-3-3ε binding defective mutant S99A led to premature disengagement, as illustrated by cells containing an increased number of pericentrin foci with just one centrin dot (Fig5G and EVFig5F). These results suggest that 14-3-3ε inhibits centrosome disengagement by suppressing Plk1 activity.

**Figure 5.**
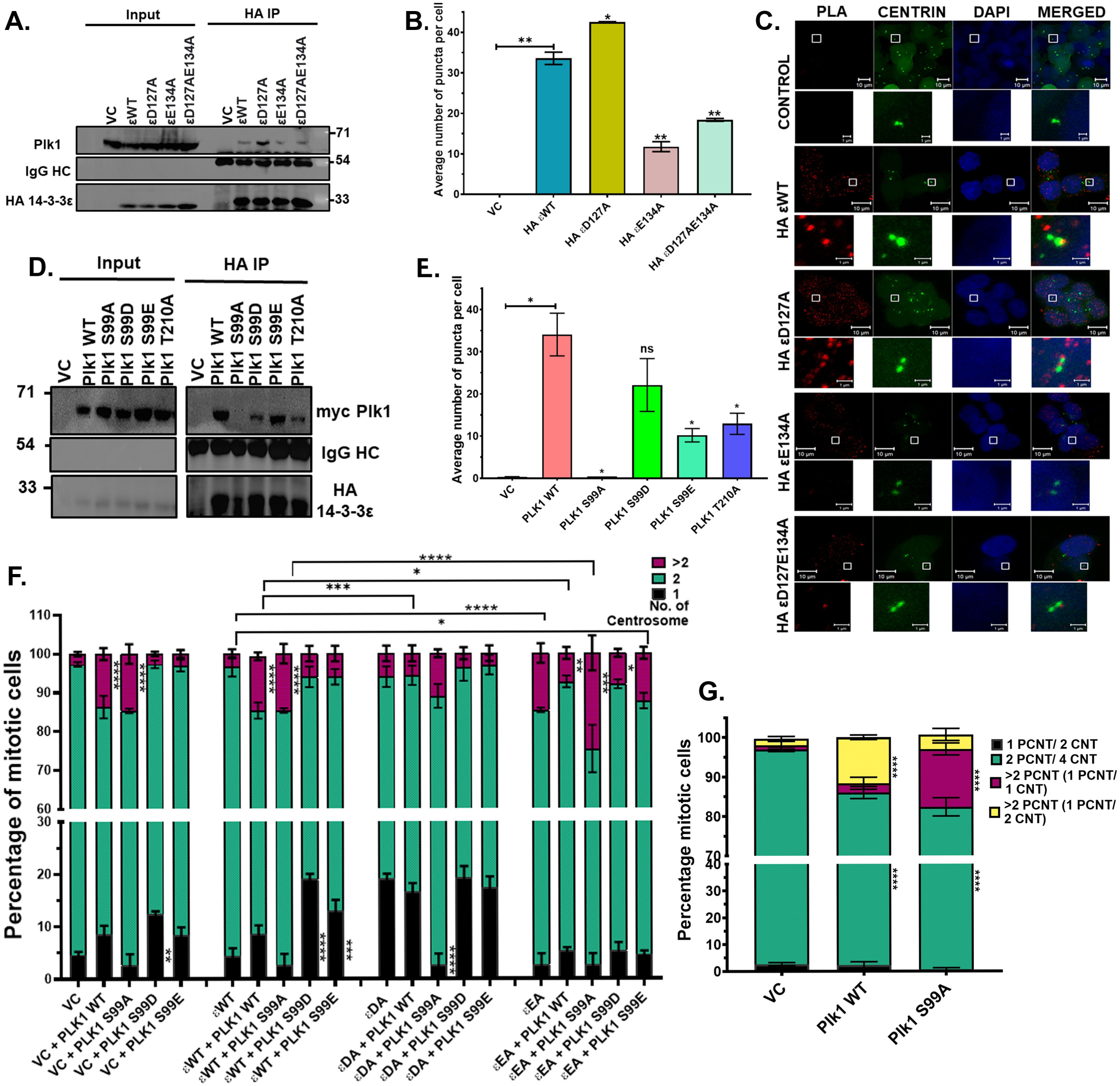
14-3-3ε inhibits Plk1, preventing centriole disengagement. **(A)** Protein extracts prepared from HCT116 cells were transfected with vector control, HA-tagged 14-3-3ε WT mutant constructs (D127A, E134A and D127AE134A) were incubated with antibodies against HA, and the reactions resolved on SDS-PAGE gels followed by Western blotting with the indicated antibodies. Input = 10%. **(B-C)** HCT116 cells stably expressing GFP Centrin2 were transfected with vector control, HA 14-3-3ε WT 14-3-3ε mutant constructs (D127A, E134A and D127AE134A) and PLA assays performed using antibodies against HA and Plk1. Representative images are shown in (C) (PLA (red), DAPI (blue)) and the mean and standard deviation of the number of puncta per cell (B). **(D-E)** HCT 116 cells expressing HA-tagged 14-3-3ε WT were co-transfected with GFP-Centrin2 and myc-tagged WT Plk1 mutant constructs (S99A, S99D, S99E and T210A). Protein extracts were prepared from these cells, and immunoprecipitations were performed with antibodies to HA, followed by Western blotting with the indicated antibodies (D). PLAs using antibodies against the HA and myc epitopes were performed, and the mean and standard deviation of the number of puncta per cell were plotted (E). **(F)** HCT116 cells stably expressing vector control and HA-tagged 14-3-3ε (WT, D127A, E134A) constructs were transfected with myc-tagged Plk1 WT and mutant constructs (S99A, S99D and S99E) followed by staining with antibodies to against the myc epitope-tag and pericentrin. The centrosome number in mitotic cells was determined in three independent experiments, and the mean and standard deviation were plotted. **(G)** HCT116 cells stably expressing the vector control, myc-tagged Plk1 WT and the Plk1 S99A mutant were stained with antibodies to the myc-epitope tag, Centrin2 and pericentrin and the PCNT/CNT ratio was determined in three independent experiments, and the mean and standard deviation plotted. p-values were obtained using unpaired student’s t-test with Welch’s correction (B and E) obtained using 2way ANOVA (Tukey’s multiple comparison) (G). *p <0.05, **p <0.01, ***p <0.001, ****p <0.0001. Scale bar = 10μm, inset = 1μm.

As 14-3-3ε forms a complex with Separase and treatment with Sepin-1 phenocopies the effect of the D127A mutant (Fig 4), we determined the ability of the WT and mutant 14-3-3ε constructs to form a complex with Separase. Co-immunoprecipitation assays demonstrated that the 14-3-3ε D127A mutant formed a complex with Separase with greater efficiency than WT 14-3-3ε (Fig6A). In contrast, the E134A mutant failed to form a complex with Separase (Fig6A). Similar results were observed in PLA assays (Fig6B and EVFig6A-B). To identify the 14-3-3ε binding site in Separase, we generated deletion mutants of Separase (Fig 6C) and tested their ability to bind 14-3-3ε in PLA assays. WT Separase and a mutant containing just the autocatalytic domain (amino acids 1141-1650) were in close proximity to 14-3-3ε, while the N-terminal and C-terminal domains of Separase (LD and AD) did not form a stable complex with 14-3-3ε (Fig6D and EVFig6C-D). Using the 14-3-3 Pred software, we identified two potential 14-3-3 binding sites in the AC domain, a Threonine at position 1363 (T1363) and a Serine at 1501 (S1501), altered them to Alanine in full-length Separase (T1363A and S1501A) and tested their ability to bind to 14-3-3ε in PLA and GST pull-down assays. The PLA assays demonstrated that T1363A formed a complex with 14-3-3ε at levels similar to WT Separase. However, the S1501A mutant failed to bind to 14-3-3ε in these assays (Fig6E and EVFig6E-F). Similar results were obtained in the GST pull-down assays (Fig6F). Next, we determined the effect of expressing the Separase mutants on the centrosome number. Expression of WT Separase or the T1363A mutant led to a significant increase in the number of cells with >2 centrosomes when co-transfected with WT 14-3-3ε or the D127A and E134A mutants of 14-3-3ε (Fig6G and EVFig6G). Expression of the S1501A mutant led to a significant increase in the number of cells with >2 centrosomes and a huge decrease in the number of cells with a single centrosome in cells expressing the D127A mutant of 14-3-3ε (Fig6G and EVFig6G). When we examined the organization of centrosomes in the cells transfected with the different Separase mutants, we observed that the cells with >2 centrosomes had just one centrin focus in one pericentrin focus and that this number was greatly elevated in cells transfected with the S1501A Separase mutant (Fig6H and EVFig6H). Finally, co-expression of the WT and/or the 14-3-3ε binding defective mutants of Separase and Plk1led to a significant increase in the number of cells with >2 centrosomes (Fig6I and EVFig6I). Expression of either the Plk1 or Separase mutant led to an increase in the number of cells with >2 centrosomes that had just one centrin focus in one pericentrin focus. This number was greatly elevated in cells transfected with both mutant constructs, suggesting a cumulative effect of the expression of both mutant proteins (Fig 6I Fig EV 6I). These results demonstrate that 14-3-3ε regulates the activity of both Plk1 and Separase to inhibit premature centriole disengagement in the centrosome duplication cycle.

**Figure 6.**
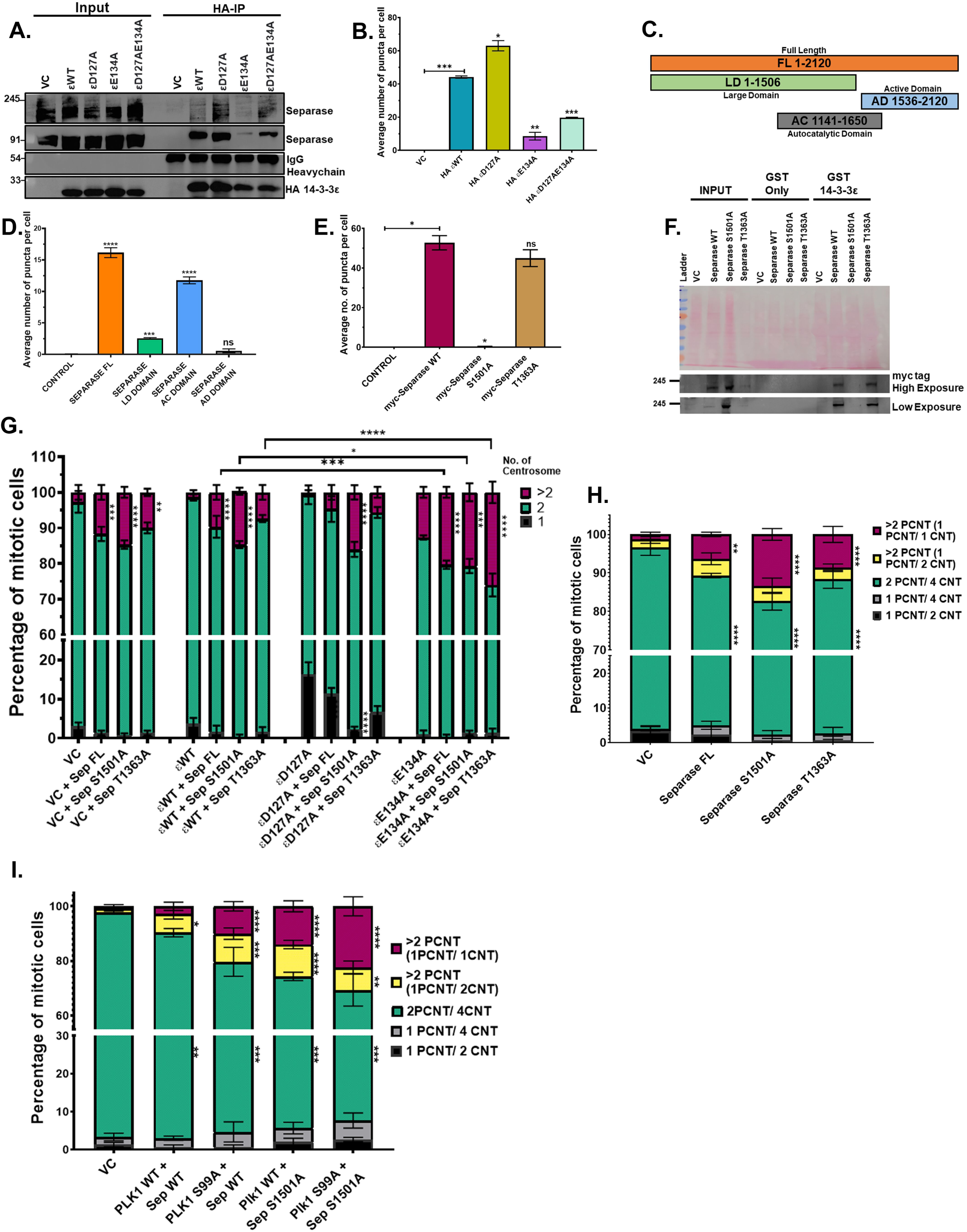
14-3-3ε inhibits Separase function, inhibiting centriole disengagement. **(A)** Protein extracts prepared from HCT116 cells transfected with vector control, HA-tagged 14-3-3ε WT mutant constructs (D127A, E134A and D127AE134A) were incubated with antibodies to HA, and the reactions resolved on SDS-PAGE gels followed by Western blotting with the indicated antibodies. **(B)** HCT116 cells stably expressing GFP Centrin2 were transfected with vector control, HA-tagged 14-3-3ε WT and mutant constructs (D127A, E134A and D127AE134A) and PLA assays performed using antibodies against HA and Separase. The number of PLA puncta per cell was determined in three independent experiments, and the mean and standard deviation were plotted. **(C)** Cartoon of the Separase deletion mutants. **(D)** Protein extracts were prepared from HCT116 cells transfected with myc-tagged Separase (FL) and myc-tagged deletion mutants (Large domain (LD), Auto-catalytic domain (AC), and Active domain (AD)). PLA assays were performed with antibodies to myc and 14-3-3ε, and the number of PLA puncta per cell was determined in three independent experiments, and the mean and standard deviation were plotted. **(E)** HCT116 cells stably expressing GFP Centrin2 were transfected with myc-tagged Separase WT and mutant constructs (S1501A and T1363A), and PLA assays were performed with antibodies to the myc epitope tag and 14-3-3ε. The number of PLA puncta per cell was determined in three independent experiments, and the mean and standard deviation were plotted. **(F)** Protein extracts prepared from HCT116 cells were transfected with myc tagged Separase WT mutant constructs (S1501A and T1363A) were incubated with the indicated GST fusion proteins followed by Western blotting with the indicated antibodies. Input = 10%. **(G)** HCT116 cells stably expressing vector control and HA-tagged 14-3-3ε (WT, D127A, E134A) constructs were transfected with myc-tagged Separase WT, and myc-tagged Separase (S1501A and T1363A) constructs, the cells arrested in mitosis followed by staining with antibodies to myc and pericentrin. The mean and standard deviation of the centrosome number for three independent experiments is plotted. **(H)** HCT116 cells stably expressing GFP Centrin2 were transfected with myc-tagged Separase WT and mutant constructs (S1501A and T1363A) and stained with antibodies against the myc epitope tag, pericentrin and the PCNT/CNT ratio was determined was determined in three independent experiments. The mean and standard deviation are plotted. **(I)** HCT116 cells stably expressing myc-tagged Plk1 WT and myc-tagged Plk1 mutants S99A were transfected with myc-tagged Separase WT, and myc-tagged Separase S1501A mutant were stained with antibodies against the myc epitope tag, pericentrin and Centrin2 and the PCNT/CNT ratio determined in three independent experiments and the mean and standard deviation plotted. p-values were obtained using 2-way ANOVA (Tukey’s multiple comparison). *p <0.05, **p <0.01, ***p <0.001, ****p <0.0001.

## DISCUSSION

The results in this report indicate that 14-3-3ε prevents premature centriole disengagement by inhibiting the function of both Plk1 and Separase. Increased complex formation between 14-3-3ε and Plk1 or Separase leads to defects in disengagement post-mitosis, thereby preventing centrosome licensing centriole duplication and monopolar spindle formation, resulting in cell death. A decrease in complex formation between 14-3-3ε and Plk1 or Separase leads to premature cleavage of the S-M linker in interphase or early mitosis, resulting in single centrioles and multi-polar mitoses, resulting in cell death. These results suggest that specific disruption of the interaction between 14-3-3ε and Plk1 or Separase could inhibit tumour progression.

Multiple reports have suggested that the loss of specific 14-3-3 paralogs leads to different phenotypes in cell lines (Dalal *et al*., 2004; Rietscher *et al*, 2018; Telles *et al*, 2009; Tilwani *et al*., 2021; Valente *et al*, 2012) and in animal models (Cheah *et al*, 2012; Sehgal *et al*., 2014; Steinacker *et al*, 2005; Tilwani *et al*, 2024; Toyo-oka *et al*., 2003; Toyo-oka *et al*, 2014). Further, structural features shared by 14-3-3ε and 14-3-3γ regulate complex formation with the mitotic phosphatase cdc25C (Telles *et al*., 2009). Recent proteomic studies that wished to identify paralog-specific interactors suggested that other than 14-3-3σ, the other 14-3-3 paralogs bound to a similar set of ligands but with different affinities, suggesting that the differences between different 14-3-3 paralogs was their affinity for the ligand rather than specific association with the ligand (Segal *et al*., 2023). However, these studies were confounded by the fact that unlike 14-3-3σ, which exclusively forms homodimers (Wilker *et al*, 2005), all the other paralogs form both homodimers and heterodimers (Yaffe *et al*., 1997; Yang *et al*, 2006b), and therefore, the low-affinity interactions might be because the target ligand might be binding to another paralog in the dimer. As 14-3-3 ligand complexes are potential drug targets, small molecules that promote disruption or formation of these complexes could serve as novel therapeutics. However, before small molecules that disrupt specific complex formation are designed, it is essential that ligands that associate specifically with individual paralogs are identified. The results in this report and our previous publications (Bose *et al*., 2021) demonstrate that the mutants that alter the conserved Aspartic acid and Glutamic acid residues to Alanine in the phospho-peptide binding permit the identification of ligands specific for each 14-3-3 paralog thereby allowing the development of these complexes as potential therapeutic targets.

The results in this report suggest that 14-3-3ε inhibits centrosome function by inhibiting the activity of Plk1 and Separase. This is consistent with previous data suggesting that Plk1 and Separase cooperate to promote centriole disengagement in human cells (Tsou *et al*., 2009). Further evidence suggests that phosphorylation of pericentrin by Plk1 results in the increased cleavage of pericentrin by Separase leading to disengagement (Kim *et al*, 2015). In addition to the cleavage of pericentrin, Plk1 also promotes the cleavage of Cep68 in prometaphase, which is followed by the degradation of pericentrin and the release of Cep215 from the centrosome leading to disengagement and licensing (Pagan *et al*., 2015). These results are consistent with our data, which illustrates that expression of the 14-3-3ε D127A mutant results in an increase in centrosome size, a decrease in the distance between centrioles and increased retention of Cep215 at the centriole. In contrast, expression of the 14-3-3ε E134A mutant results in a decrease in centrosome size, increased distance between centrioles and decreased retention of Cep215 at the centriole. Further, treatment with Plk1 or Separase inhibitors exacerbates the phenotypes observed with the D127A mutant and prevents premature centriole disengagement in interphase observed in cells expressing the E134A mutant. In contrast, expression of a Plk1 or Separase mutants that do not bind to 14-3-3ε leads to increased disengagement and centriole splitting in cells expressing the E134A mutant. Further, while treatment with both a Plk1 inhibitor and a Separase inhibitor prevents premature centriole disengagement, treatment with both inhibitors does not significantly decrease centriole disengagement. However, co-expression of mutants in Plk1 and Separase that do not bind to 14-3-3ε results in a significant increase in centriole disengagement and premature centriole cleavage, a result distinct from that observed for the dual inhibitor treatment. These results could be because Plk1 is upstream of Separase in the disengagement pathway and suggest that 14-3-3ε inhibits the activity of both Plk1 and Separase to prevent premature disengagement.

In addition to the cleavage of the cohesion complex leading to sister chromatid separation at Anaphase, Separase is also required for centriole disengagement by stimulating the cleavage of pericentrin (Lee & Rhee, 2012; Matsuo *et al*, 2012). Multiple reports have suggested that cleavage of the cohesion complex is synchronized with centriole disengagement to permit the timely completion of mitosis (Nakamura *et al*, 2009; Schockel *et al*., 2011; Thein *et al*, 2007), though other reports have suggested that disengagement might occur before sister chromatid separation (Agircan & Schiebel, 2014). However, all of these reports suggest that premature Separase activation can lead to premature disengagement and centriole splitting (Karki *et al*., 2017; Nakamura *et al*., 2009; Schockel *et al*., 2011; Thein *et al*., 2007). These results are consistent with our observations that expression of the 14-3-3ε E134A mutant, which does not bind effectively to Separase, results in centriole disengagement and premature centriole splitting in interphase cells.

In addition to regulating centriole disengagement at the end of mitosis, both Plk1 and Separase are required for other functions during mitosis. Plk1-mediated phosphorylation of Cdc6 leads to complex formation between Cdc6 and cyclin B/cdk1, leading to Separase activation (Yim & Erikson, 2010). In addition, Plk1 phosphorylates cdc25C leading to a cdc25C nuclear transport, activating the cyclinB/cdk1 complex and mitotic progression (Toyoshima-Morimoto *et al*, 2002). As 14-3-3ε is required to inhibit cdc25C function in interphase, preventing the premature activation of the cyclin B/cdk1 complex (Dalal *et al*., 2004), it is possible that in addition to binding to and inhibiting Plk1 and Separase activity, loss of 14-3-3ε or the expression of the E134A mutant of 14-3-3ε, could indirectly lead to an increase in Plk1 and Separase activity leading to premature centriole disengagement. This is consistent with our observation that the 14-3-3ε E134A mutant shows decreased complex formation with cdc25C and previous reports demonstrating that 14-3-3ε inhibits cdc25C function (Dalal *et al*., 2004; Telles *et al*., 2009). However, treatment with a cdk1 inhibitor does not lead to a decrease in the phenotypes associated with the expression of the E134A mutant, suggesting that the effects of 14-3-3ε on centriole disengagement might be independent of the ability of 14-3-3ε to inhibit cdc25C function.

Previous results have demonstrated that Plk1 forms a complex with 14-3-3γ and 14-3-3ζ (Du *et al*., 2012; Kasahara *et al*., 2013). 14-3-3ζ binds to Plk1 phosphorylated at S330 and S597; these residues are distinct from S99, and the 14-3-3ζ Plk1 interaction regulates cytokinesis (Du *et al*., 2012). Further, the loss of 14-3-3ζ does not lead to centrosome amplification (this report), unlike the loss of 14-3-3ε (Mukhopadhyay *et al*., 2016; Tilwani *et al*., 2021). 14-3-3γ binds to Plk1 when it is phosphorylated at S99 (Kasahara *et al*., 2013), the same residue that we have identified as required for complex formation with 14-3-3ε (this report) and loss of either 14-3-3γ or 14-3-3ε leads to centrosome amplification (Mukhopadhyay *et al*., 2016; Tilwani *et al*., 2021). However, our previous results suggest that 14-3-3γ inhibits centriole duplication by binding to NPM1 and that expression of the 14-3-3γ Aspartic acid mutant leads to the formation of a disengaged centrosome that cannot induce centriole duplication (Bose *et al*., 2021), a phenotype distinct from that reported for 14-3-3ε in this report. These results suggest that the two 14-3-3 paralogs regulate distinct steps in the centrosome duplication pathway.

Previous reports in the literature have demonstrated that 14-3-3ε and 14-3-3γ are present in centrosomal fractions (Pietromonaco *et al*., 1996). Likewise, the loss of these paralogs leads to centrosome amplification in multiple human cell lines and in mice (Mukhopadhyay *et al*., 2016; Tilwani *et al*., 2024; Tilwani *et al*., 2021), unlike the loss of 14-3-3ζ (this report). cdk1 activation is first observed at the centrosome (Jackman *et al*, 2003) and the Nek2 pathway, which is required for centrosome disjunction, is activated by the cyclin B2/cdk1 complex (Nam & van Deursen, 2014). The cyclin B/cdk1 complex is activated by the mitotic phosphatase cdc25C (Lee *et al*, 1992; O’Connor *et al*, 1994; Strausfeld *et al*, 1991), and the activity of this phosphatase is inhibited by 14-3-3ε and 14-3-3γ (Dalal *et al*., 1999; Dalal *et al*., 2004; Hosing *et al*, 2008; Telles *et al*., 2009), which activates the cyclin B/cdk1 complex. The premature activation of the cyclin B/cdk1 complex upon loss of either 14-3-3ε or 14-3-3γ leads to centrosome amplification (Mukhopadhyay *et al*., 2016). Centrosome amplification is a feature of multiple tumor types show (Bose & Dalal, 2019b) and while loss of only 14-3-3γ leads to tumour progression, loss of both 14-3-3ε and 14-3-3γ can lead to increased cell death in vitro and tumour regression in vivo (Mukhopadhyay *et al*., 2016). This is consistent with data showing that while loss of both paralogs results in centrosome amplification, cells with a loss of 14-3-3ε show multipolar mitoses leading to a decrease in proliferation and survival and loss of 14-3-3γ leads to clustered mitoses leading to increased survival (Tilwani *et al*., 2021). The loss of 14-3-3γ disrupts desmosome formation, leading to decreased cell stiffness and increased frequency of clustered mitoses and is consistent with other data from the literature showing that increased cell-cell adhesion prevents clustered mitoses and promotes multi-polar mitoses (Rhys *et al*, 2018), which is associated with decreased survival in tumor cells (Ganem *et al*., 2009). Therefore, targeting complex formation between 14-3-3ε and Plk1 or Separase, leading to premature disengagement and multi-polar mitoses, could be an excellent approach for killing tumor cells.

Our results suggest that the expression of the D127A and E134A mutants leads to the formation of monopolar spindles and multi-polar spindles, decreasing proliferation and cell death. Similarly, loss of 14-3-3ε in the mouse epidermis results in a decrease in proliferation and a decrease in the thickness of the mouse epidermis, which correlates with an increase in centrosome number in the basal layer of the epidermis in these mice (Tilwani *et al*., 2024). Similar results have been reported when Plk4 is over-expressed in the basal layer of the epidermis (Kulukian *et al*, 2015) suggesting that centrosome amplification and multi-polar spindle formation might lead to defects in the orientation of the mitotic spindle in the basal layer thereby inhibiting the asymmetric divisions required for epidermal differentiation.

The results in this report lead to the following model (Fig7). 14-3-3ε inhibits the activity of Plk1 and Separase in late interphase and part of mitosis, preventing premature centriole disengagement. The 14-3-3ε D127A mutant binds with greater affinity to both Plk1 and Separase, inhibiting centriole disengagement through interphase. This leads to monopolar spindle formation in mitosis, resulting in cell death. In contrast, expression of the E134A mutant that shows a decreased affinity for Plk1 and Separase leads to premature centriole disengagement and centriole splitting, resulting in the formation of multipolar spindles, which post cytokinesis results in cell death. In conjunction with our previous studies, these results demonstrate that different 14-3-3 paralogs regulate different stages in the centrosome cycle and argue for paralog-specific regulation of a subset of protein ligands.

**Figure 7.**
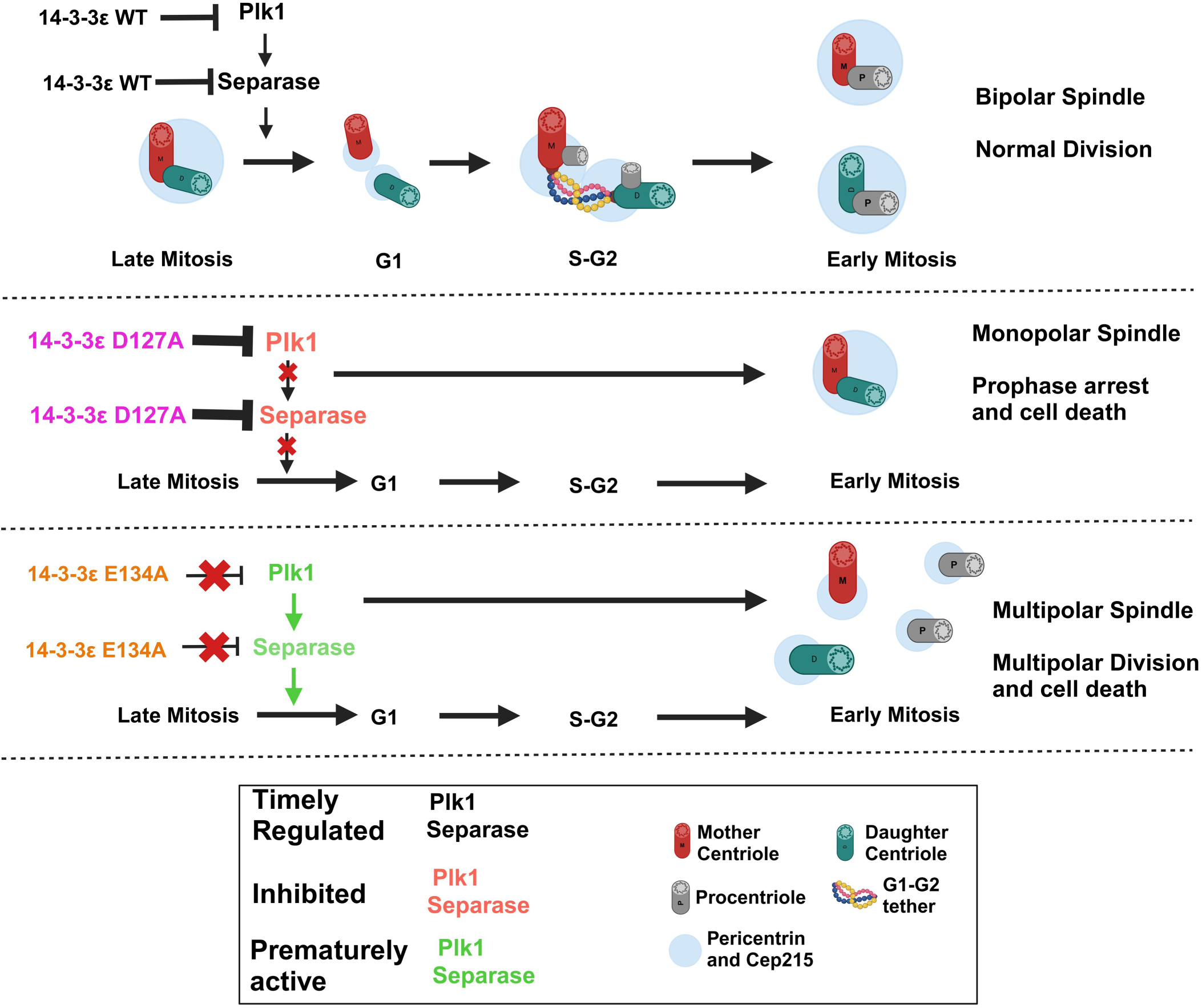
Model for regulation of centriole disengagement by 14-3-3ε. 14-3-3ε inhibits premature centriole disengagement by inhibiting the activity of Plk1 and Separase. 14-3-3ε mutants showing increased or decreased affinity for Plk1 and Separase inhibit disengagement or promote premature centriole disengagement, respectively. Image created using Biorender.

## MATERIALS AND METHODS

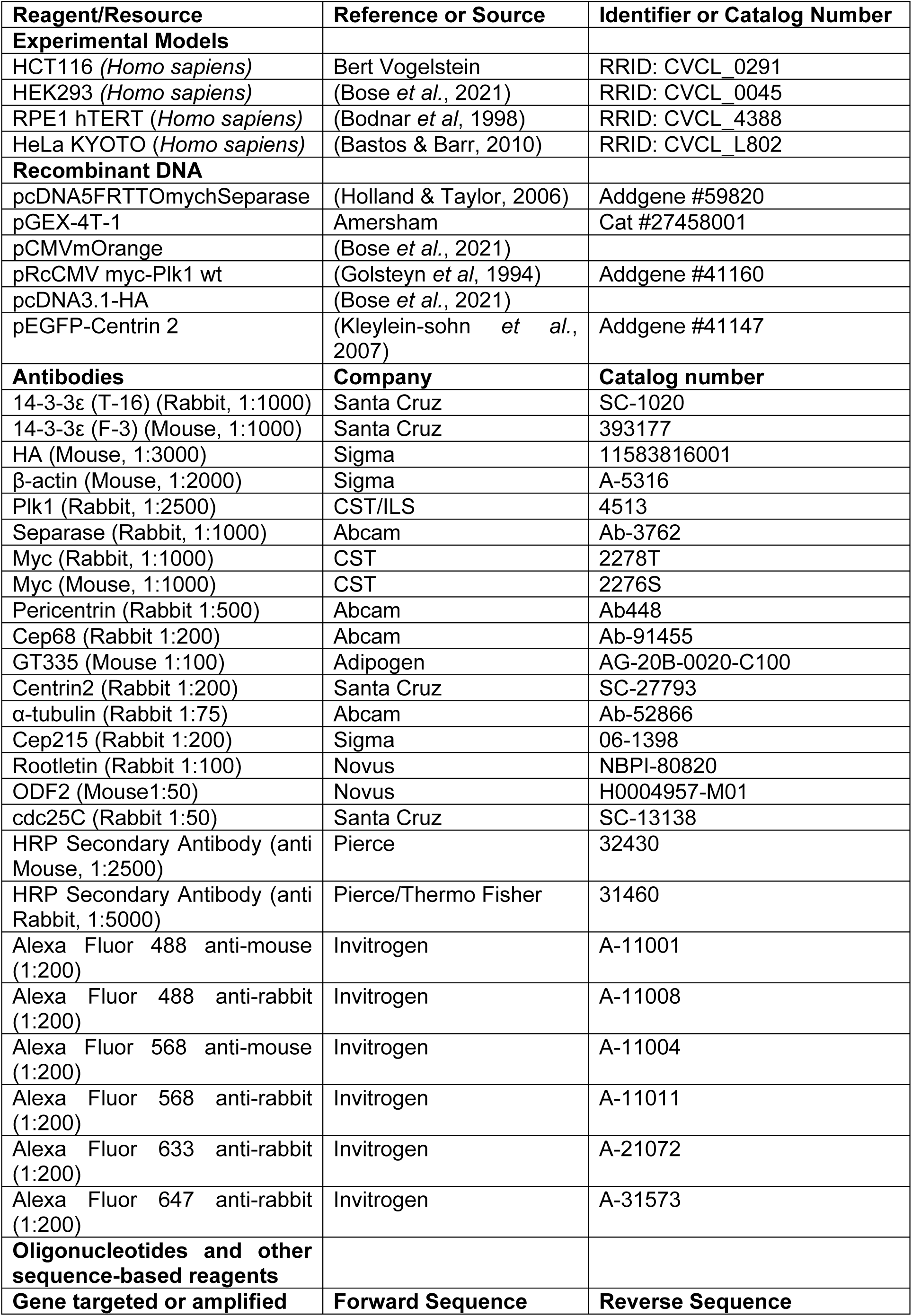

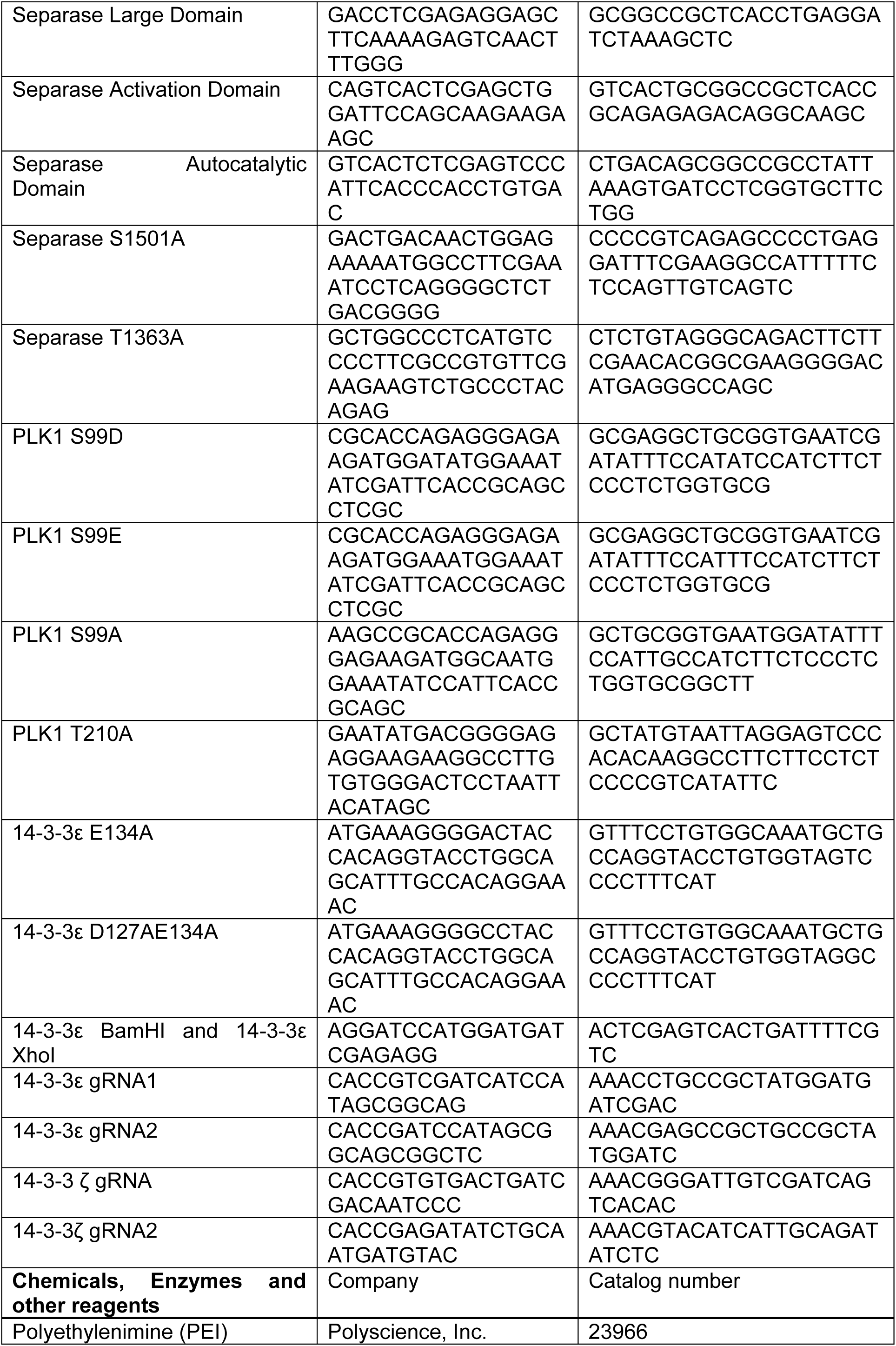

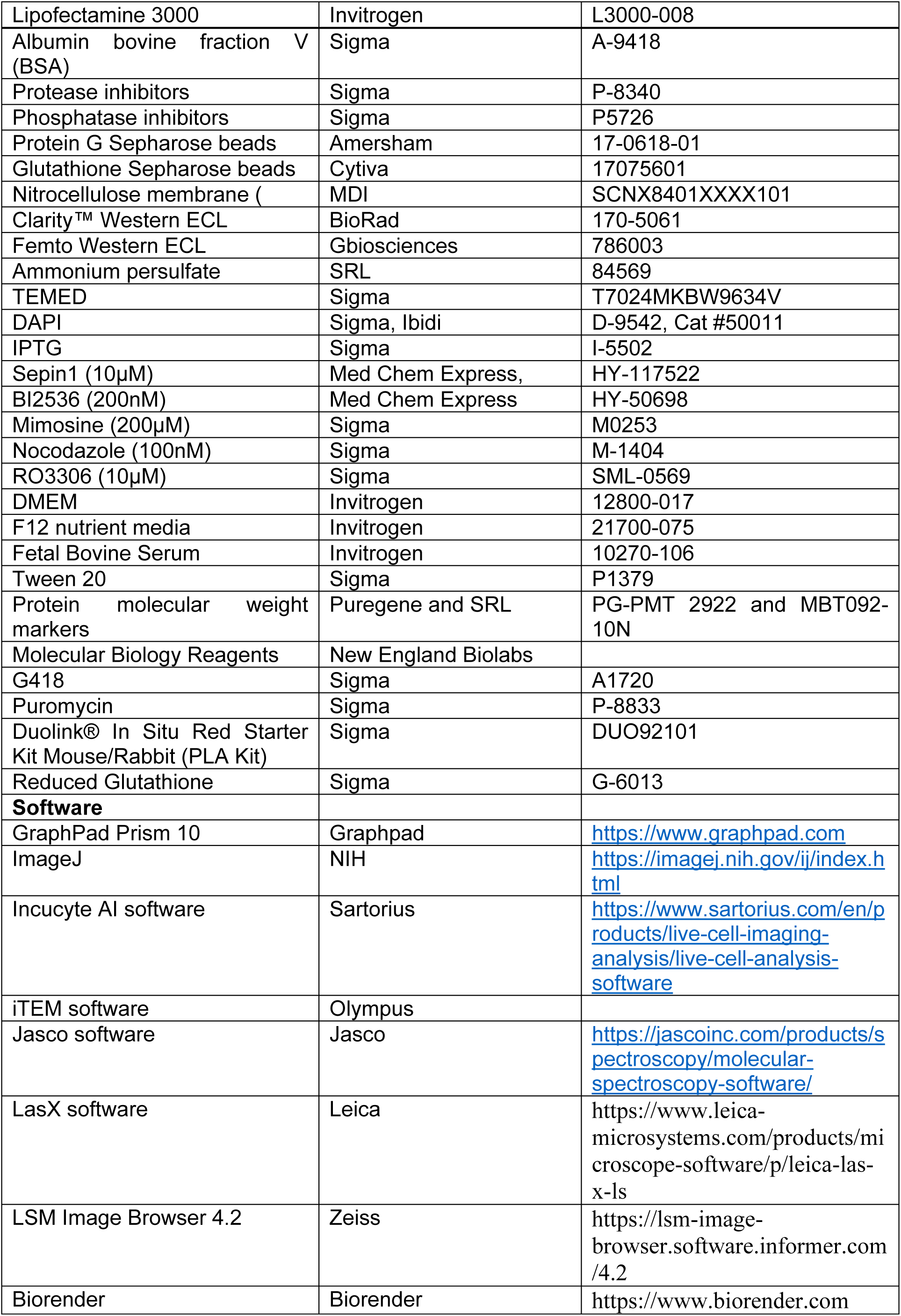

### Cell lines and transfections

HCT116 (RRID: CVCL_0291) HEK293 (RRID: CVCL_0045), hTERT-RPE1 (RRID: CVCL_4388) and HeLa Kyoto EGFP-alpha-tubulin/H2B-mCherry (RRID: CVCL_L802) cells were cultured as described previously (Bose *et al*., 2021; Tilwani *et al*., 2021). All the cell lines were verified by short tandem repeat (STR) profiling and were mycoplasma free. HCT116 cells derived 14-3-3ε and 14-3-3ζ knockout lines were generated as described (Tilwani *et al*., 2021) using the guide RNA sequences in table 1. Transfections were performed with Polyethylenimine (PEI) (Polysciences, Inc.) or Lipofectamine 3000 (Invitrogen) as per manufacturer’s instructions. Where indicated cells were treated with the vehicle control or the following drugs at the indicated concentrations for 20-24 hours followed by immunofluorescence analysis Western blotting, and flow cytometry. Drug concentrations are in the table.

### Plasmids

The 14-3-3ε derived mutant constructs (D127A, E134A and D127AE134A) were generated by site directed mutagenesis as described (Vishal *et al*, 2018). WT and mutant 14-3-3ε WT constructs were cloned into pCMVmOrange digested with EcoRI and XhoI. The HA tagged 14-3-3ε WT and mutants (D127A, E134A and D127AE134A) were amplified in polymerase chain reactions and cloned into pCDNA3HA (Bose *et al*., 2021) digested with BamHI and XhoI. pcDNA5 FRT TO myc hSeparase was a gift from Stephen Taylor (Holland & Taylor, 2006) (Addgene plasmid number #59820) the mutants (LD, AD, AC, S1501A and T1363A) were generated using site directed mutagenesis or amplified in polymerase chain reactions. pRcCMV myc-Plk1 WT (Addgene plasmid #41160), (Addgene plasmid number #41158) (Golsteyn *et al*., 1994) and pEGFP-Centrin2 (Addgene plasmid #41147) (Kleylein-sohn *et al*., 2007) were gifts from Erich Nigg and the Plk1 point mutants (S99D, S99E, S99A, and T210A) were generated site directed mutagenesis. The14-3-3ε WT and mutants (D127A, E134Aand D127AE134A) were cloned into pGEXT-4T-1 using primers shown in table 1. All oligonucleotide sequences are listed in the reagent table.

### Western blot analysis

Protein extracts were prepared and resolved on SDS-PAGE gels followed by Western blotting with antibodies to indicated proteins in table 2 as previously described (Bose *et al*., 2021) as described and proteins were transferred to nitrocellulose membrane and probed. The blots were developed with Clarity (Bio-Rad) Western blot chemiluminescent substrate (Clarity™ Western ECL Substrate), Femto (Bio-Rad) Western blot chemiluminescent substrate or Femtolucent plus HRP chemiluminescent reagents and imaged on a ChemiDoc™ Imaging System (Bio-Rad).

### Immunofluorescence assays

Immunofluorescence Assays (IFA) were performed as previously described (Bose *et al*., 2021). Briefly, post-transfection, the cells were synchronized or treated with inhibitors as described above and fixed and stained with the indicated antibodies (Reagent table). Confocal images were acquired on an LSM 780 Carl Zeiss confocal microscope or Leica SP8 confocal microscope at a magnification of 630x or 1000x with 5x-6x digital zoom. An image of the entire cell was acquired with 0.35μm sections. Acquired images were processed using LSM Image Browser or LASX software, where the image was represented in 2-D by the projection of the entire z-stacks. The insets were derived from the Extract Region tool in the LSM Image Browser. The quantitation of intensity of Cep 215, pericentrin the area of centrosome and the distance between the centriole was calculated in a minimum of 30 cells per set in three independent experiments using Image J FIJI software. For determining the number of centrosomes, PCNT/CNT ratio, Rootletin/CNT, Cep68/CNT and ODF2/CNT, 100 cells per set expressing the desired constructs were manually counted either on an LSM 780 Carl Zeiss confocal microscope or Leica SP8 confocal microscope or Olympus Inverted IX73 fluorescence or Zeiss AXIO Imager.Z1 upright fluorescence microscope in three independent experiments.

### Proximity Ligation Assays

HCT116 cells were cultured in 8 well chambered slide at the confluency of about 50%, transfected with the respective constructs and fixed with 4%PFA (Paraformaldehyde) for 20 mins and then PLA was performed using Duolink® PLA Reagent by following Duolink® PLA Fluorescence protocol. Briefly, fixed cells were permeabilize (using 0.3% Triton X-100 and 0.1%NP-40), blocked (using Blocking solution) and then incubated with the respective primary antibody dilution (using Antibody Diluent) mentioned in the table 1 for either 2 hours at RT or overnight at 4°C. The cells were washed thrice with 1x wash buffer 1 and incubated with PLA Probes (PLUS and MINUS) followed by Duolink® Fluorescence Detection Reagent (red or far red) which includes ligation and amplification. The cells were washed twice with 1x wash buffer post ligation and amplification followed by a final wash with 0.01x Wash Buffer B for 1 min. Cells were then mounted using 10μl of mounting medium with DAPI and images were acquired using an LSM 780 Carl Zeiss confocal microscope at a magnification of 400x 0r 630x with 4-6x optical zoom and images processed as described above. Z-stack projected images were analyzed using the Image J FIJI software to determine the average number of PLA puncta per cell. Total number of puncta (red) per field was divided by the total number of cells per field (DAPI-blue) to get the average number of puncta per cell. The mean and standard deviation of three independent experiments is plotted.

### Electron Microscopy

HCT116 cells stably expressing Vector Control, and HA tagged 14-3-3ε WT and mutants (D127A and E134A) were synchronized in S-G2 with RO3306 for 18h, trypsinized, and processed as described (Tilwani *et al*., 2021). The grids were imaged on JEM 1400 PLUS, transmission electron microscope. Images were analyzed using iTEM software (OSIS,GmbH) and Imaje J Fiji software to calculate the centriole length, inner diameter and outer diameter. Minimum number of cells analyzed to determine the length of the centriole was N ≥ 15 and for determining inner and outer diameter N ≥ 30 across three sets.

### Live Cell Imaging, proliferation assays and flow cytometry

HeLa KYOTO cells were transfected with HA pcDNA3 vector control, HA-tagged 14-3-3ε WT, and HA-tagged 14-3-3ε (D127A and E134A), and stable cells were generated using G418. Post selection, the expression was confirmed by Western blotting, and the cells were subjected to live cell imaging on Olympus 3i Spinning Disc. For the inhibitor experiment, the cells were treated with vehicle (DMSO), 200nM BI2536 or 10μM Sepin1 or both and the images were acquired at 600x magnification using 488 and 568 excitation wavelengths, in the following condition: 37°C, 5% CO2, and 75% humidity. The images of the same cell were acquired at an interval of 15-20 mins over 20-24h. The video and images were processed using Slide Book 6 software, showing the time stamp at the top and scale at the bottom.

HCT116 cells cultured in glass bottom 24 well plates were transfected with vector control, mOrange tagged 14-3-3ε WT and mutant (D127A and E134A) constructs. 24h post-transfection, the proliferation was measured by acquiring the images using 568 excitation wavelength and phase contrast of the entire well at intervals of 1h over 48h, for each construct, on Sartorius Incucyte Live cell imaging system using 20X objective. The cells were maintained at 37°C, 5% CO2, and 70% humidity during acquisition. The data was analyzed by training the Incucyte AI software by selecting random fields to sense the number of mOrange positive cells and the total number of cells (using phase contrast). The fold change was determined by dividing the total number of mOrange positive cells per field by initial number of mOrange positive cells (phase contrast) per field for each time point and plotted.

Trypsinized HCT116 cells treated with the drugs were washed with PBS, and fixed with ice cold 70% Ethanol and then incubated with 0.1mg/ml Propidium Iodide and RNase for 30 mins and the samples were run on Thermo Fisher Attune NxT Flow Cytometer and data analysis and graph were obtained using Modfit software.

### GST pulldown and co-immunoprecipitation assays

GST pulldown and co-immunoprecipitation assays were performed as described previously (Bose *et al*., 2021). HCT116 cells were grown up to 70% confluency and were transfected with either myc tag Plk1 (WT, S99A, S99D, S99E and T210A) constructs or myc tagged hSeparase (WT, S1501A and T1363A) constructs. 48 hours post-transfection, cells were lysed in EBC lysis buffer (0.5mM Tris pH 8.0, 125 mM NaCl, 0.5% (v/v) NP-40, protease inhibitor cocktail, phosphatase inhibitor cocktail). Following centrifugation, 10% input was removed and lysates were incubated with equal amount of GST only, 14-3-3εWT and 14-3-3ε mutants (D127A, 14-3-3εE134A, 14-3-3εD127AE134A) GST fusion proteins bound to glutathione beads for either 4-5h at RT or 16h at 4°C. Post incubation, 3x washes with 50mM NET-N (1M Tris pH 8.0, 2.5M NaCl, 0.5M EDTA pH 8.0, 0.5% (v/v) NP-40) were given post which the complexes were resolved on 10% SDS-PAGE gels followed by Western blotting.

For endogenous immunoprecipitation, near confluent HCT116 cells were lysed in EBC buffer, 10% input was removed and the remaining supernatant was divided into two, one was incubated with 14-3-3ε or Plk1 or Separase the other with IgG Rabbit. Post 3 hrs of incubation, 50% slurry of protein G-Sepharose beads was added for 1h at 4°C and then 3x washes with 50mM NET-N was given. The complex was resolved on 10% SDS-PAGE gels followed by Western blotting.

For co-immunoprecipitation, 70% confluent HCT116 cells were transfected with HA vector control, HA tagged 14-3-3ε (WT, D127A, E134A and D127AE134A) along with myc tag Plk1 (WT, S99A, S99D, S99E and T210A) constructs. Lysates were prepared and incubated for 3h at 4°C with HA antibody(12CA5) to precipitate HA-tagged proteins. The reaction was incubated with protein G-Sepharose beads for 1h at 4°C and washed thrice with 50mM NET-N (1M Tris pH 8.0, 2.5M NaCl, 0.5M EDTA pH 8.0, 0.5% (v/v) NP-40). The reactions were resolved on 10% SDS-PAGE gels followed by Western blotting.

### CD spectra

pGEX-4T-1 14-3-3ε WT and mutant constructs were purified as described (Bose *et al*., 2021). The recombinant proteins were eluted with 10mM reduced glutathione and dialysed against with 20mM Tris pH8, NaCl 20mM overnight. The proteins were concentrated to 15-20 μM and CD performed on Jasco J-1500 CD Spectrophotometer and the data processed in Jasco software.

## ACKNOWLEDGEMENTS.

The authors wish to acknowledge the ACTREC Digital Imaging Facility, the Flow Cytometry Facility, Biophysics Facility and TEM Facility for help with image acquisition and we thank E. Nigg for the GFP-Centrin2 construct. The experiments in this report were funded by a grant from the Department of Biotechnology (BT/PR38272/BRB/10/1894/2020), Department of Atomic Energy DPR Nos. 1/3(7)/2020/TMC/R&D-II/ 8823 and 1/3(6)/2020/TMC/R&D -II/ 3805 and donations to the ACTREC Basic Research fund (4338) to SND

## Author contributions

Monika A. Jaiswal: Data curation; Investigation; Methodology; Visualization; Writing—review and editing. Akshay Karn: Methodology. Aparna Das: Methodology. Anisha Kumari: Methodology. Shilu Tiwari: Methodology. Sorab N. Dalal: Funding acquisition; Supervision; Writing—review and editing.

## Data availability

All relevant data can be found within the article and its supplementary information.

## Expanded View Figure Legends

**Expanded View Figure 1. 14-3-3ε mutants alter centrosome numbers in different cell lines. (A-C)** Representative images of the indicated cell lines transfected with the mOrange tagged wild type (WT) and mutant (D127A, E134A, D127AE134A)14-3-3ε constructs. The quantitation is in Figure 1C-E. **(D)** Western blot analysis showing equal expression of the different 14-3-3ε constructs in the different cell lines. β-actin served as a loading control. **(E-H)** HCT116 (E) and RPE1 Tert (F) derived 14-3-3ε knockout lines transfected with the indicated constructs were stained with antibodies to pericentrin, and representative images are shown. The quantitation is in Figure 1F-G. Protein extracts from the transfected cells were resolved on SDS-PAGE gels, followed by Western blotting with the indicated antibodies. β-actin served as a loading control. **(I-K)** HCT116 derived 14-3-3ζ knockout cells were generated. Protein extracts prepared from the vector control and 14-3-3ζ knockout cells were resolved on SDS PAGE gels, followed by Western blotting with the indicated antibodies (I). Note that the knockout cells have low levels of 14-3-3ζ. The vector control and knockout cells were arrested in mitosis, stained with antibodies to pericentrin, and counterstained with DAPI. Representative images are shown (J), and the mean and standard deviation of three independent experiments are plotted (K). p-values were obtained using unpaired student’s t-test with Welch’s correction. ns – non-significant, *p < 0.05. Scale= 10µm.

**Expanded View Figure 2. 14-3-3ε prevents centriole disengagement. (A)** Western blot showing the expression of mOrange tagged 14-3-3ε constructs (WT, D127A, E134A, D127AE134A) for experiments shown in figure 2B-D. β-actin served as a loading control. **(B-C)** Representative images for the experiments are shown in Figure 2D (B) and Figure 2H (C). **(D)** Western blot showing the expression of mOrange tagged 14-3-3ε constructs (WT, D127A, E134A, D127AE134A) for experiments shown in figure 2F-J and EV figure 2C. β-actin served as a loading control. Scale bar = 10μm, inset scale bar = 1μm.

**Expanded View Figure 3. Expression of the 14-3-3ε mutants leads to a mitotic delay and cell death. (A)** Representative blot shows the ectopic expression of mOrange tagged, and HA-tagged 14-3-3ε constructs (WT, D127A, E134A and D127AE134A) in HCT116 cells and HeLa KYOTO for experiments shown in figure 3. β-actin served as a loading control. **(B)** HCT116 cells transfected with the indicated constructs were stained with antibodies to glutamylated tubulin (GT335 green) and counter-stained with DAPI (blue). Scale = 5μm, inset scale = 0.5μm. **(C)** Electron micrographs showing centriole structure indicated with a white arrow in cells stably expressing HA vector control, and HA-tagged 14-3-3ε constructs (WT, D127A, EI134A and DI27AE134A). Scale= 2μm, 500nm, 200nm. **(D)** HCT116 cells expressing mOrange vector control and mOrange tagged 14-3-3ε constructs (WT, D127A, E134A, D127AE134A) were synchronized in mitosis using nocodazole followed by washing out of the nocodazole allowing mitotic progression. Cells were fixed and stained for Pericentrin (green), α-tubulin (red) and counter-stained with DAPI (blue). Scale= 5μm.

**Expanded View Figure 4. 14-3-3ε forms a complex with Plk1 and Separase. (A)** Ponceau image of the GST pulldown blot showing band for GST only and GST 14-3-3ε WT and GST 14-3-3ε K50E. **(B-C)** Protein extracts from the HCT116 cells were incubated with antibodies to Separase (B) or Plk1 (C), and the reactions resolved on SDS PAGE gels followed by Western blotting with the indicated antibodies. Immunoprecipitations with IgG served as negative controls. **(D)** Protein extracts prepared from HCT116 cells transfected with the indicated constructs and treated with BI2536 or Sepin-1 were resolved on SDS PAGE gels, followed by Western blotting with the indicated antibodies. β-actin served as a loading control. **(E)** The cell cycle distribution of the indicated cells treated with either BI2536 or Sepin-1. The vehicle control, mimosine and nocodazole-treated cells served as controls. **(F-H)** HCT116 cells stably expressing GFP Centrin2 were transfected with mOrange vector control and mOrange tagged 14-3-3ε constructs (WT, D127A, E134A) and treated with BI2536, Sepin1 or both (BI2536 and Sepin1) and stained for pericentrin and counterstained with DAPI. Representative images of cells treated with BI2536 (F) with Sepin-1 are shown in (G), and both BI2536 and Sepin-1 (H) are shown. **(I-K)** HeLa KYOTO cells stably expressing HA 14-3-3ε WT or HA 14-3-3ε E134A were treated with vehicle control (DMSO), BI2536, Sepin-1 or both BI2536 and Sepin-1 and were subjected to live cell imaging at intervals of 15 mins. Representative images of the same cells treated with BI2536 (I), Sepin-1 (J) or both BI2536 and Sepin1(K) are shown, and the time stamp is present in the top left corner and the scale is represented in the bottom right. Scale = 10μm.

**Expanded View Figure 5. 14-3-3ε inhibits Plk1, preventing centriole disengagement. (A)** Protein extracts prepared from HCT116 cells were incubated with GST alone, or GST-tagged WT and mutant (D127A, E134A and D127AE134A) 14-3-3ε fusion proteins immobilized on glutathione Sepharose beads. The reactions were resolved on SDS PAGE gels, and Western blots were performed with the indicated antibodies. GST only served as a negative control, and cdc25C served as a positive control to study the interaction of 14-3-3ε with Plk1. **(B)** A Circular Dichroism (CD) analysis was performed to determine the secondary structure of WT and mutant (D127A and E134A) 14-3-3ε GST fusion proteins. The spectra for the individual proteins are shown. **(C-D)** HCT116 cells stably expressing the vector control or myc-tagged WT and mutant (S99A, S99D, S99E and T210A) Plk1 constructs were to perform PLA assays. Representative images are shown (C), and a Western blot demonstrates that the Plk1 proteins were present at equivalent levels (D). β-actin served as a loading control. **(E)** HCT116 cells stably expressing vector control and HA-tagged 14-3-3ε (WT, D127A and E134A) constructs were transfected with myc-tagged WT and mutant (S99A, S99D and S99E) Plk1constructs, and immunofluorescence staining was performed using antibodies to the myc-epitope tag and pericentrin and the representative images are shown (myc-tag (orange), pericentrin (red) and DAPI (blue)). **(F)** HCT116 cells stably expressing myc-tagged Plk1 WT and Plk1 S99A were arrested in mitosis and stained with antibodies to Centrin2 and pericentrin and counterstained with DAPI. Representative images are shown. Scale = 10μm and = 1µm in the inset.

**Expanded View Figure 6. 14-3-3ε inhibits Separase function, inhibiting centriole disengagement. (A-B)** HCT116 cells stably expressing GFP Centrin2 were transfected with the vector control or HA-tagged WT and mutant (D127A, E134A and D127AE134A) 14-3-3ε constructs and PLA assays were performed. A Western blot analysis demonstrates that the constructs were expressed at similar levels (A). Representative images are shown in (B). Scale = 10μm and inset scale = 0.5μm. **(C-D)** HCT116 cells were transfected with different domains of myc-tagged full-length Separase (FL) and myc-tagged deletion mutants of Separase (Large domain (LD), Auto-catalytic domain (AC), and Active domain (AD)) and PLA assays performed with antibodies to 14-3-3ε and the myc-epitope tag. Representative images are shown in (C). Scale = 20μm and inset scale = 5μm. All Separase mutant constructs were expressed at equivalent levels, and Western blots for β-actin served as loading controls (D). **(E-F)** HCT116 cells stably expressing GFP Centrin2 were transfected with myc-tagged WT and mutant (S1501A and T1363A) Separase constructs and PLA assays performed with antibodies to 14-3-3ε and the myc-epitope tag. Representative images are shown in (E), and a Western blot analysis demonstrates that the proteins were present at similar levels (F). Scale = 20μm and inset scale = 2μm. **(G)** HCT116 cells stably expressing vector control and HA-tagged 14-3-3ε (WT, D127A, E134A) constructs were transfected with myc-tagged Separase WT and myc-tagged Separase (S1501A and T1363A) constructs and immunofluorescence staining was performed using antibodies against myc and pericentrin. Representative images are shown in (G). Scale = 10μm. **(H)** HCT116 cells stably expressing GFP Centrin2 were transfected with myc-tagged Separase WT and myc-tagged Separase mutants (S1501A and T1363A) and stained with antibodies against myc (orange) and pericentrin (red) and counterstained with DAPI (blue). Representative images are shown (H). Scale = 10μm and inset scale = 1μm. **(I)** HCT116 cells stably expressing myc-tagged Plk1 WT and myc-tagged Plk1 mutants S99A were transfected with myc-tagged Separase WT and myc-tagged Separase S1501A and stained with antibodies against pericentrin (red) and Centrin2 (green) and counterstained with DAPI (blue). Scale = 10μm and inset scale = 1μm.

## Expanded View Videos

**Video 1-4. Mitotic phenotypes observed upon expression of WT and mutant 14-3-3ε.** HeLa Kyoto cells stably expressing the vector control (1) or WT 14-3-3ε (2), or the 14-3-3ε mutants D127A (3) or E134A (4) were imaged over time on an Olympus 3i spinning disc microscope at intervals of 20 minutes over 20 hours. The time stamp is on the top right, and the scale bar is on the bottom left. Scale bar = 10µm.

**Video 5-12. Mitotic phenotypes observed upon treatment of cells expressing WT and mutant 14-3-3ε with Plk1 and Separase inhibitors.** HeLa Kyoto cells stably expressing WT 14-3-3ε (5-8), or the 14-3-3ε E134A mutant (9-12) were treated with the vehicle control (5,9), the Plk1 inhibitor BI2536 (6,10), the Separase inhibitor Sepin-1 (7,11) or both inhibitors (8,12). The cells were imaged over time on an Olympus 3i spinning disc microscope at intervals of 20 minutes over 20 hours. The time stamp is on the top right, and the scale bar is on the bottom left. Scale bar = 10µm.

## BIBLIOGRAPHY

Abal M, Keryer G, Bornens M (2005) Centrioles resist forces applied on centrosomes during G2/M transition. Biol Cell 97: 425–434

Adon AM, Zeng X, Harrison MK, Sannem S, Kiyokawa H, Kaldis P, Saavedra HI (2010) Cdk2 and Cdk4 regulate the centrosome cycle and are critical mediators of centrosome amplification in p53-null cells. Mol Cell Biol 30: 694–710

Agircan FG, Schiebel E (2014) Sensors at centrosomes reveal determinants of local separase activity. PLoS Genet 10: e1004672

Aitken A (2006) 14-3-3 proteins: a historic overview. Semin Cancer Biol 16: 162–172

Andersen JS, Wilkinson CJ, Mayor T, Mortensen P, Nigg EA, Mann M (2003) Proteomic characterization of the human centrosome by protein correlation profiling. Nature 426: 570–574

Bahe S, Stierhof YD, Wilkinson CJ, Leiss F, Nigg EA (2005) Rootletin forms centriole-associated filaments and functions in centrosome cohesion. J Cell Biol 171: 27–33

Basto R, Brunk K, Vinadogrova T, Peel N, Franz A, Khodjakov A, Raff JW (2008) Centrosome Amplification Can Initiate Tumorigenesis in Flies. Cell Reports 133: 11

Bastos RN, Barr FA (2010) Plk1 negatively regulates Cep55 recruitment to the midbody to ensure orderly abscission. J Cell Biol 191: 751–760

Bobinnec Y, Khodjakov A, Mir LM, Rieder CL, Edde B, Bornens M (1998a) Centriole disassembly in vivo and its effect on centrosome structure and function in vertebrate cells. J Cell Biol 143: 1575–1589

Bobinnec Y, Moudjou M, Fouquet JP, Desbruyeres E, Edde B, Bornens M (1998b) Glutamylation of centriole and cytoplasmic tubulin in proliferating non-neuronal cells. Cell Motil Cytoskeleton 39: 223–232

Bodnar AG, Ouellette M, Frolkis M, Holt SE, Chiu CP, Morin GB, Harley CB, Shay JW, Lichtsteiner S, Wright WE (1998) Extension of life-span by introduction of telomerase into normal human cells. Science 279: 349–352

Bornens M (2002) Centrosome composition and microtubule anchoring mechanisms. Curr Opin Cell Biol 14: 25–34

Bose A, Dalal SN (2019a) 14-3-3 proteins mediate the localization of Centrin2 to the centrosome. J Biosciences 44

Bose A, Dalal SN (2019b) Centrosome Amplification and Tumorigenesis: Cause or Effect? Results Probl Cell Differ 67: 413–440

Bose A, Modi K, Dey S, Dalvi S, Nadkarni P, Sudarshan M, Kundu TK, Venkatraman P, Dalal SN (2021) 14-3-3gamma prevents centrosome duplication by inhibiting NPM1 function. Genes to cells : devoted to molecular & cellular mechanisms 26: 426–446

Brinkley BR, Cox SM, Pepper DA, Wible L, Brenner SL, Pardue RL (1981) Tubulin assembly sites and the organization of cytoplasmic microtubules in cultured mammalian cells. J Cell Biol 90: 554–562

Brunet A, Kanai F, Stehn J, Xu J, Sarbassova D, Frangioni JV, Dalal SN, DeCaprio JA, Greenberg ME, Yaffe MB (2002) 14-3-3 transits to the nucleus and participates in dynamic nucleo-cytoplasmic transport. J Cell Biol 156: 817–828

Castellanos E, Dominguez P, Gonzalez C (2008) Centrosome dysfunction in Drosophila neural stem cells causes tumors that are not due to genome instability. Curr Biol 18: 1209–1214

Cheah PS, Ramshaw HS, Thomas PQ, Toyo-Oka K, Xu X, Martin S, Coyle P, Guthridge MA, Stomski F, van den Buuse M et al (2012) Neurodevelopmental and neuropsychiatric behaviour defects arise from 14-3-3zeta deficiency. Mol Psychiatry 17: 451–466

Coblitz B, Shikano S, Wu M, Gabelli SB, Cockrell LM, Spieker M, Hanyu Y, Fu H, Amzel LM, Li M (2005) C-terminal recognition by 14-3-3 proteins for surface expression of membrane receptors. J Biol Chem 280: 36263–36272

Dalal SN, Schweitzer CM, Gan J, DeCaprio JA (1999) Cytoplasmic localization of human cdc25C during interphase requires an intact 14-3-3 binding site. Mol Cell Biol 19: 4465–4479

Dalal SN, Yaffe MB, DeCaprio JA (2004) 14-3-3 family members act coordinately to regulate mitotic progression. Cell Cycle 3: 672–677

Du J, Chen L, Luo X, Shen Y, Dou Z, Shen J, Cheng L, Chen Y, Li C, Wang H et al (2012) 14-3-3zeta cooperates with phosphorylated Plk1 and is required for correct cytokinesis. Front Biosci (Schol Ed) 4: 639–650

Fry AM (2002) The Nek2 protein kinase: a novel regulator of centrosome structure. Oncogene 21: 6184–6194

Ganem NJ, Godinho SA, Pellman D (2009) A mechanism linking extra centrosomes to chromosomal instability. Nature 460: 278–282

Golsteyn RM, Schultz SJ, Bartek J, Ziemiecki A, Ried T, Nigg EA (1994) Cell cycle analysis and chromosomal localization of human Plk1, a putative homologue of the mitotic kinases Drosophila polo and Saccharomyces cerevisiae Cdc5. J Cell Sci 107 (Pt 6): 1509–1517

Graser S, Stierhof YD, Nigg EA (2007) Cep68 and Cep215 (Cdk5rap2) are required for centrosome cohesion. J Cell Sci 120: 4321–4331

Habedanck R, Stierhof YD, Wilkinson CJ, Nigg EA (2005) The Polo kinase Plk4 functions in centriole duplication. Nat Cell Biol 7: 1140–1146

Heald R, Khodjakov A (2015) Thirty years of search and capture: The complex simplicity of mitotic spindle assembly. J Cell Biol 211: 1103–1111

Hinchcliffe EH, Sluder G (2001) “It takes two to tango”: understanding how centrosome duplication is regulated throughout the cell cycle. Genes Dev 15: 1167–1181

Holland AJ, Taylor SS (2006) Cyclin-B1-mediated inhibition of excess separase is required for timely chromosome disjunction. J Cell Sci 119: 3325–3336

Hosing AS, Kundu ST, Dalal SN (2008) 14-3-3 Gamma is required to enforce both the incomplete S phase and G2 DNA damage checkpoints. Cell Cycle 7: 3171–3179

Jackman M, Lindon C, Nigg EA, Pines J (2003) Active cyclin B1-cdk1 first appears on centrosomes in prophase. Nat Cell Biol 5: 143–148

Karki M, Keyhaninejad N, Shuster CB (2017) Precocious centriole disengagement and centrosome fragmentation induced by mitotic delay. Nature communications 8: 15803

Kasahara K, Goto H, Izawa I, Kiyono T, Watanabe N, Elowe S, Nigg EA, Inagaki M (2013) PI 3-kinase-dependent phosphorylation of Plk1-Ser99 promotes association with 14-3-3gamma and is required for metaphase-anaphase transition. Nature communications 4: 1882

Kim J, Lee K, Rhee K (2015) PLK1 regulation of PCNT cleavage ensures fidelity of centriole separation during mitotic exit. Nature communications 6: 10076

Kleylein-sohn J, Westendorf J, Clech ML, Habedanck R, Stierhof Y-d, Nigg EA (2007) Plk4-Induced Centriole Biogenesis in Human Cells. Developmental Cell: 23

Kulukian A, Holland AJ, Vitre B, Naik S, Cleveland DW, Fuchs E (2015) Epidermal development, growth control, and homeostasis in the face of centrosome amplification. Proc Natl Acad Sci U S A 112: E6311–6320

Kumagai A, Dunphy WG (1999) Binding of 14-3-3 proteins and nuclear export control the intracellular localization of the mitotic inducer Cdc25. Genes & development 13: 1067–1072

Lee K, Rhee K (2012) Separase-dependent cleavage of pericentrin B is necessary and sufficient for centriole disengagement during mitosis. Cell Cycle 11: 2476–2485

Lee MS, Ogg S, Xu M, Parker LL, Donoghue DJ, Maller JL, Piwnica-Worms H (1992) cdc25+ encodes a protein phosphatase that dephosphorylates p34cdc2. Mol Biol Cell 3: 73–84

Lenart P, Petronczki M, Steegmaier M, Di Fiore B, Lipp JJ, Hoffmann M, Rettig WJ, Kraut N, Peters JM (2007) The small-molecule inhibitor BI 2536 reveals novel insights into mitotic roles of polo-like kinase 1. Curr Biol 17: 304–315

Levine MS, Bakker B, Boeckx B, Moyett J, Lu J, Vitre B, Spierings DC, Lansdorp PM, Cleveland DW, Lambrechts D et al (2017) Centrosome Amplification Is Sufficient to Promote Spontaneous Tumorigenesis in Mammals. Dev Cell 40: 313–322.e315

Liu D, Bienkowska J, Petosa C, Collier RJ, Fu H, Liddington R (1995) Crystal structure of the zeta isoform of the 14-3-3 protein. Nature 376: 191–194

Mahen R, Venkitaraman AR (2012) Pattern formation in centrosome assembly. Curr Opin Cell Biol 24: 14–23

Mardin BR, Agircan FG, Lange C, Schiebel E (2011) Plk1 controls the Nek2A-PP1gamma antagonism in centrosome disjunction. Curr Biol 21: 1145–1151

Mardin BR, Lange C, Baxter JE, Hardy T, Scholz SR, Fry AM, Schiebel E (2010) Components of the Hippo pathway cooperate with Nek2 kinase to regulate centrosome disjunction. Nat Cell Biol 12: 1166–1176

Matsuo K, Ohsumi K, Iwabuchi M, Kawamata T, Ono Y, Takahashi M (2012) Kendrin is a novel substrate for separase involved in the licensing of centriole duplication. Curr Biol 22: 915–921

Meraldi P, Lukas J, Fry AM, Bartek J, Nigg EA (1999) Centrosome duplication in mammalian somatic cells requires E2F and Cdk2-cyclin A. Nat Cell Biol 1: 88–93

Modi K, Dalvi S, Venkatraman P (2020) Two negatively charged invariant residues influence ligand binding and conformational dynamics of 14-3-3zeta. FEBS Lett 594: 878–886

Moorhead G, Douglas P, Morrice N, Scarabel M, Aitken A, MacKintosh C (1996) Phosphorylated nitrate reductase from spinach leaves is inhibited by 14-3-3 proteins and activated by fusicoccin. Curr Biol 6: 1104–1113

Mukhopadhyay A, Sehgal L, Bose A, Gulvady A, Senapati P, Thorat R, Basu S, Bhatt K, Hosing AS, Balyan R et al (2016) 14-3-3gamma Prevents Centrosome Amplification and Neoplastic Progression. Sci Rep 6: 26580

Muslin AJ, Tanner JW, Allen PM, Shaw AS (1996) Interaction of 14-3-3 with signaling proteins is mediated by recognition of phosphoserine. Cell 84: 889–897

Nakamura A, Arai H, Fujita N (2009) Centrosomal Aki1 and cohesin function in separase-regulated centriole disengagement. J Cell Biol 187: 607–614

Nam HJ, van Deursen JM (2014) Cyclin B2 and p53 control proper timing of centrosome separation. Nat Cell Biol 16: 538–549

Nigg EA, Stearns T (2011) The centrosome cycle: Centriole biogenesis, duplication and inherent asymmetries. Nat Cell Biol 13: 1154–1160

O’Connor PM, Ferris DK, Hoffmann I, Jackman J, Draetta G, Kohm KW (1994) Role of the cdc25C phosphatase in G2 arrest induced by nitrogen mustard. Proc Natl Acad Sci USA 91: 9480–9484

Obsil T, Ghirlando R, Klein DC, Ganguly S, Dyda F (2001) Crystal structure of the 14-3-3zeta:serotonin N-acetyltransferase complex. a role for scaffolding in enzyme regulation. Cell 105: 257–267

Ohta M, Watanabe K, Ashikawa T, Nozaki Y, Yoshiba S, Kimura A, Kitagawa D (2018) Bimodal Binding of STIL to Plk4 Controls Proper Centriole Copy Number. Cell Reports 23: 3160–3169.e3164

Pagan JK, Marzio A, Jones MJ, Saraf A, Jallepalli PV, Florens L, Washburn MP, Pagano M (2015) Degradation of Cep68 and PCNT cleavage mediate Cep215 removal from the PCM to allow centriole separation, disengagement and licensing. Nat Cell Biol 17: 31–43

Peng CY, Graves PR, Thoma RS, Wu Z, Shaw AS, Piwnica-Worms H (1997) Mitotic and G2 checkpoint control: regulation of 14-3-3 protein binding by phosphorylation of Cdc25C on serine-216. Science 277: 1501–1505

Pietromonaco SF, Seluja GA, Aitken A, Elias L (1996) Association of 14-3-3 proteins with centrosomes. Blood Cells Mol Dis 22: 225–237

Pihan GA (2013) Centrosome dysfunction contributes to chromosome instability, chromoanagenesis, and genome reprograming in cancer. Frontiers in oncology 3: 277

Reber S, Hyman AA (2015) Emergent Properties of the Metaphase Spindle. Cold Spring Harb Perspect Biol 7: a015784

Rhys AD, Monteiro P, Smith C, Vaghela M, Arnandis T, Kato T, Leitinger B, Sahai E, McAinsh A, Charras G et al (2018) Loss of E-cadherin provides tolerance to centrosome amplification in epithelial cancer cells. J Cell Biol 217: 195–209

Rietscher K, Keil R, Jordan A, Hatzfeld M (2018) 14-3-3 proteins regulate desmosomal adhesion via plakophilins. J Cell Sci 131

Rittinger K, Budman J, Xu J, Volinia S, Cantley LC, Smerdon SJ, Gamblin SJ, Yaffe MB (1999) Structural analysis of 14-3-3 phosphopeptide complexes identifies a dual role for the nuclear export signal of 14-3-3 in ligand binding. Mol Cell 4: 153–166

Schockel L, Mockel M, Mayer B, Boos D, Stemmann O (2011) Cleavage of cohesin rings coordinates the separation of centrioles and chromatids. Nat Cell Biol 13: 966–972

Segal D, Maier S, Mastromarco GJ, Qian WW, Nabeel-Shah S, Lee H, Moore G, Lacoste J, Larsen B, Lin ZY et al (2023) A central chaperone-like role for 14-3-3 proteins in human cells. Mol Cell 83: 974–993 e915

Sehgal L, Mukhopadhyay A, Rajan A, Khapare N, Sawant M, Vishal SS, Bhatt K, Ambatipudi S, Antao N, Alam H (2014) 14-3-3γ-Mediated transport of plakoglobin to the cell border is required for the initiation of desmosome assembly in vitro and in vivo. Journal of cell science 127: 2174–2188

Shen YH, Godlewski J, Bronisz A, Zhu J, Comb MJ, Avruch J, Tzivion G (2003) Significance of 14-3-3 self-dimerization for phosphorylation-dependent target binding. Mol Biol Cell 14: 4721–4733

Shukla A, Kong D, Sharma M, Magidson V, Loncarek J (2015) Plk1 relieves centriole block to reduplication by promoting daughter centriole maturation. Nature communications 6: 8077

Sir JH, Putz M, Daly O, Morrison CG, Dunning M, Kilmartin JV, Gergely F (2013) Loss of centrioles causes chromosomal instability in vertebrate somatic cells. J Cell Biol 203: 747–756

Steegmaier M, Hoffmann M, Baum A, Lenart P, Petronczki M, Krssak M, Gurtler U, Garin-Chesa P, Lieb S, Quant J et al (2007) BI 2536, a potent and selective inhibitor of polo-like kinase 1, inhibits tumor growth in vivo. Curr Biol 17: 316–322

Steinacker P, Schwarz P, Reim K, Brechlin P, Jahn O, Kratzin H, Aitken A, Wiltfang J, Aguzzi A, Bahn E et al (2005) Unchanged survival rates of 14-3-3gamma knockout mice after inoculation with pathological prion protein. Mol Cell Biol 25: 1339–1346

Strausfeld U, Labbe JC, Fesquet D, Cavadore JC, Picard A, Sadhu K, Russell P, Doree M (1991) Dephosphorylation and activation of a p34^cdc2^/cyclin B complex in vitro by human cdc25 protein. Nature 351: 242–245

Sullenberger C, Vasquez-Limeta A, Kong D, Loncarek J (2020) With Age Comes Maturity: Biochemical and Structural Transformation of a Human Centriole in the Making. Cells 9

Telles E, Hosing AS, Kundu ST, Venkatraman P, Dalal SN (2009) A novel pocket in 14-3-3e is required to mediate specific complex formation with cdc25C and to inhibit cell cycle progression upon activation of checkpoint pathways. Exp Cell Res 315: 1448–1457

Thein KH, Kleylein-Sohn J, Nigg EA, Gruneberg U (2007) Astrin is required for the maintenance of sister chromatid cohesion and centrosome integrity. J Cell Biol 178: 345–354

Tilwani S, Gandhi K, Dalal SN (2024) 14-3-3epsilon conditional knockout mice exhibit defects in the development of the epidermis. FEBS Lett

Tilwani S, Gandhi K, Narayan S, Ainavarapu SRK, Dalal SN (2021) Disruption of desmosome function leads to increased centrosome clustering in 14-3-3gamma-knockout cells with supernumerary centrosomes. FEBS Lett 595: 2675–2690

Toyo-oka K, Shionoya A, Gambello MJ, Cardoso C, Leventer R, Ward HL, Ayala R, Tsai LH, Dobyns W, Ledbetter D et al (2003) 14-3-3epsilon is important for neuronal migration by binding to NUDEL: a molecular explanation for Miller-Dieker syndrome. Nat Genet 34: 274–285

Toyo-oka K, Wachi T, Hunt RF, Baraban SC, Taya S, Ramshaw H, Kaibuchi K, Schwarz QP, Lopez AF, Wynshaw-Boris A (2014) 14-3-3epsilon and zeta regulate neurogenesis and differentiation of neuronal progenitor cells in the developing brain. J Neurosci 34: 12168–12181

Toyoshima-Morimoto F, Taniguchi E, Nishida E (2002) Plk1 promotes nuclear translocation of human Cdc25C during prophase. EMBO Rep 3: 341–348

Tsou MF, Stearns T (2006) Mechanism limiting centrosome duplication to once per cell cycle. Nature 442: 947–951

Tsou MF, Wang WJ, George KA, Uryu K, Stearns T, Jallepalli PV (2009) Polo kinase and separase regulate the mitotic licensing of centriole duplication in human cells. Dev Cell 17: 344–354

Tzivion G, Luo Z, Avruch J (1998) A dimeric 14-3-3 protein is an essential cofactor for Raf kinase activity. Nature 394: 88–92

Valente C, Turacchio G, Mariggio S, Pagliuso A, Gaibisso R, Di Tullio G, Santoro M, Formiggini F, Spano S, Piccini D et al (2012) A 14-3-3gamma dimer-based scaffold bridges CtBP1-S/BARS to PI(4)KIIIbeta to regulate post-Golgi carrier formation. Nat Cell Biol 14: 343–354

Vishal SS, Tilwani S, Dalal SN (2018) Plakoglobin localization to the cell border restores desmosome function in cells lacking 14-3-3gamma. Biochem Biophys Res Commun 495: 1998–2003

Whalley HJ, Porter AP, Diamantopoulou Z, White GR, Castaneda-Saucedo E, Malliri A (2015) Cdk1 phosphorylates the Rac activator Tiam1 to activate centrosomal Pak and promote mitotic spindle formation. Nature communications 6: 7437

Wilker EW, Grant RA, Artim SC, Yaffe MB (2005) A structural basis for 14-3-3sigma functional specificity. J Biol Chem 280: 18891–18898

Wong YL, Anzola JV, Davis RL, Yoon M, Motamedi A, Kroll A, Seo CP, Hsia JE, Kim SK, Mitchell JW et al (2015) Cell biology. Reversible centriole depletion with an inhibitor of Polo-like kinase 4. Science 348: 1155–1160

Xiao B, Smerdon SJ, Jones DH, Dodson GG, Soneji Y, Aitken A, Gamblin SJ (1995) Structure of a 14-3-3 protein and implications for coordination of multiple signalling pathways. Nature 376: 188–191

Yaffe MB (2002) How do 14-3-3 proteins work?--Gatekeeper phosphorylation and the molecular anvil hypothesis. FEBS Lett 513: 53–57

Yaffe MB, Rittinger K, Volinia S, Caron PR, Aitken A, Leffers H, Gamblin SJ, Smerdon SJ, Cantley LC (1997) The structural basis for 14-3-3 phosphopeptide binding specificity. Cell 91: 961–971

Yang J, Adamian M, Li T (2006a) Rootletin interacts with C-Nap1 and may function as a physical linker between the pair of centrioles/basal bodies in cells. Mol Biol Cell 17: 1033–1040

Yang K, Tylkowski MA, Huber D, Contreras CT, Hoyer-Fender S (2018) ODF2 maintains centrosome cohesion by restricting beta-catenin accumulation. J Cell Sci 131

Yang X, Lee WH, Sobott F, Papagrigoriou E, Robinson CV, Grossmann JG, Sundstrom M, Doyle DA, Elkins JM (2006b) Structural basis for protein-protein interactions in the 14-3-3 protein family. Proc Natl Acad Sci U S A 103: 17237–17242

Yim H, Erikson RL (2010) Cell division cycle 6, a mitotic substrate of polo-like kinase 1, regulates chromosomal segregation mediated by cyclin-dependent kinase 1 and separase. Proc Natl Acad Sci U S A 107: 19742–19747

Zhang N, Scorsone K, Ge G, Kaffes CC, Dobrolecki LE, Mukherjee M, Lewis MT, Berg S, Stephan CC, Pati D (2014) Identification and Characterization of Separase Inhibitors (Sepins) for Cancer Therapy. J Biomol Screen 19: 878–889

